# Predictive encoding of auditory sequences in the human prefrontal cortex

**DOI:** 10.1101/2025.08.22.671264

**Authors:** Olgerta Asko, Vegard A. Volehaugen, Anaïs Llorens, Ingrid Funderud, Sabine L. Leske, Jugoslav Ivanovic, Torstein R. Meling, Lars Etholm, Pål Gunnar Larsson, Robert T. Knight, Tor Endestad, Anne-Kristin Solbakk, Alejandro O. Blenkmann

## Abstract

Humans extract regularities from the environment to form expectations that guide perception and optimize behavior. Although the prefrontal cortex (PFC) is central to this process, the relative contributions of orbitofrontal (OFC) and lateral PFC (LPFC) remain unclear. Here, we show that the brain tracks sound regularities in an auditory deviance detection task to predict when a target deviant will occur. Intracranial EEG in epilepsy patients reveals prefrontal engagement, with earlier expectancy-related modulation in OFC and later modulation in LPFC. Connectivity analyses indicate bidirectional and asymmetrical expectancy-related information exchange between the two areas, with a first lead by OFC, consistent with its role in initiating predictive encoding. Converging causal evidence shows that OFC lesions abolish sensitivity to expectancy, whereas LPFC lesions yield only modest effects not significantly different from controls. Together, these results provide electrophysiological and causal evidence for distinct, temporally organized contributions of prefrontal subregions to predictive processing.

**Significance Statement:** The human brain builds probabilistic associations capturing the dynamic environmental structure in a predictive manner. The ability to encode and deploy predictive information to anticipate future events relies on the prefrontal cortex (PFC). However, the specific roles of PFC subregions in this process remain unclear. Here, we provide converging electrophysiological and causal lesion evidence on the distinct involvement of the orbital and lateral PFC in generating anticipatory neural signals based on probabilistic auditory structure.

## Introduction

In everyday life, we encounter predictive sensory cues that signal a need for imminent action. These cue-action associations can be built and stored in memory through years of experience, such as stepping on the brake when the traffic light switches from green to yellow, or using rhythmic and melodic cues to prepare motions on the dance floor. This implies that the brain does not solely process sensory signals, but also infers their likely occurrence and outcomes^1–3^ to prepare subsequent action. Inferences are formatted by tracking events’ occurrence or transition probabilities^3–9^ and serve as the basis for anticipation, initiating perceptual, cognitive, and motor operations proactively.^10,11^ Learning probabilistic predictive structure can engage anticipatory neural activity, which facilitates processing of the predicted events and optimizes behavior.^12–15^

The ability to use predictive information to guide perception and action is known to rely on the prefrontal cortex (PFC), which exerts a facilitatory top-down bias on sensory pathways based on internal representations of current information and prior knowledge.^16–21^ The predictive coding framework has also implicated PFC in higher-order forming of predictions, prediction error processing, and top-down modulation of deviance detection.^22–29^ There is, however, sparse knowledge about the specific roles of PFC subregions in predictive processes.

Orbitofrontal cortex (OFC) has long been linked to inhibitory control,^30–33^ flexible learning,^34–37^ and decision-making.^38–41^ Increasing research suggests that OFC generates predictions about upcoming events and action outcomes by integrating contextual and temporal attributes^42^ to update mental representations and guide behavior.^43–49^ This aligns with the view of OFC as a key part of a neural circuit that forms, maintains, and updates a mental model of the environment or a so-called “cognitive map” of task state space,^50^ with OFC monitoring the individual’s current position within this space.^51,52^ This state-space representation holds relationships between task-relevant states (i.e., task stages or conditions), which are unobservable from current sensory input and determined only by the trajectory of previous events.^46,53–55^ This capacity allows tracking event regularities over time to build and update an internal representational (i.e., predictive) model of the external environment to predict upcoming events and action outcomes.

Although evidence for lateral PFC (LPFC) is sparser, recent fMRI work shows that while medial OFC represents hidden task-state information, lateral OFC and dorsolateral PFC encode the abstract rules governing structure knowledge across task states by integrating immediate prior information with current sensory inputs.^56^ LPFC is also involved in prospective functions supporting predictive processes, including selective attention, working memory, response preparation, and inhibition.^16,57–59^ LPFC facilitates sensory processing by enhancing and accelerating sensory responses.^60–62^ In audition, LPFC plays a key role in focusing attention on relevant sounds and filtering out distracting ones,^63^ with inferior frontal gyrus (IFG) involved in structured sequence processing,^64^ essential for detecting deviance embedded in extended stimulus dependencies.^24–26,65–69^ Specifically, the right IFG is implicated in detecting unpredicted auditory transitions,^69,70^ whereas the left IFG is particularly involved in detecting unpredicted sound durations and gaps.^24^

To differentiate the roles of the OFC and LPFC, and delineate their causal contribution in encoding auditory environmental regularities to anticipate future events, we studied a reliable neural marker of anticipation, the Contingent Negative Variation (CNV), also known as the “expectancy wave”.^71^ The CNV is a slow ramping event-related potential (ERP) arising between two contingently associated sensory events reflecting both cognitive and motor preparation.^14,72–77^ CNV generation is influenced by explicit ^78–80^ and implicit stimulus factors,^81–83^ such as probability and valence, state-related factors such as arousal and attention,^22,73,84,85^ as well as neurological conditions.^86,87^ The development of the CNV is linked to a distributed network of cortical regions, including premotor and supplementary motor, cingulo-insular, superior and inferior parietal, and superior and middle temporal areas, along with the key engagement of lateral and orbital PFC.^85,88–93^ Lesion studies have demonstrated the importance of the dorsolateral PFC (DLPFC),^94–96^ with intracranial EEG studies further implicating the OFC.^97–99^ The configuration of the neural generators is determined by multiple factors, including higher-order cognitive representations, such as task strategy, but also sensorimotor characteristics of the task, including motor output demands and sensory input modality and structure.^100,101^ The role of the CNV during multiscale sensory information delivery is rarely investigated; with only few studies exploring its modulation during exposure to hierarchically nested auditory sequences.^22,102,103^

In the local-global paradigm,^22,28^ tone sequences are structured at two levels. At the local level, a tone change or an omission may happen after four repetitive tones. At the global level, a particular type of tone sequence occurs frequently, reflecting the higher-order structure. Global deviants, which violate this rule, are infrequent and random. We focused on the global structure of the task and predicted that the CNV would evolve as the probability of encountering a deviant tone sequence increased, reflecting the brain’s ability to form implicit expectations based on global regularities. A pivotal study showed that the CNV was enhanced during deviant tone sequences occurring after several repetitions of standard sequences, reflecting the degree of expectancy of the upcoming deviating sequence.^22^ Thus, information about states in a task-space might be extracted from the temporal structure (i.e., information not present in the current sensory input) to facilitate expectation of global stimulus violations. We hypothesized that CNV modulation could index the implicit expectation about the imminence of uncertain deviant sequences that might be based on a mental representation or “cognitive map” of the overarching task structure.

In a previous paper, we reported electrophysiological support for the involvement of OFC in detecting violations of auditory predictions at two hierarchical levels of task structure.^104^ Specifically, the results indicated that OFC supports a top-down predictive process that modulates deviant events expectation in lower sensory areas. In the present study, we aimed to further examine the roles of OFC and LPFC in building global expectations based on implicit predictive information embedded in the task structure. We recorded scalp EEG activity during an auditory local-global deviance detection task in which participants counted deviant tone sequences. The recordings were obtained from patients with OFC or LPFC lesions, as well as a healthy control group. In parallel, we recorded intracranial EEG (stereoelectroencephalography; SEEG) from epilepsy patients implanted with depth electrodes in OFC or LPFC. Additionally, we conducted a behavioral study in healthy participants using a reaction time (RT) version of the task, designed to assess whether global expectations facilitate detection of deviant events by enhancing response speed.

For healthy adults, we hypothesized that the neuronal encoding of the temporal structure of non-deviant and deviant sequences would progressively increase CNV amplitude across successive sequence steps prior to the occurrence of a random deviant. We further hypothesized that the anticipatory process indexed by the CNV would also be encoded in intracerebrally recorded high-frequency broadband activity (HFBA; 65 – 145 Hz),^105,106^ which is a localized proxy for macroscale spiking activity near the recording electrode.^107–109^ Given the PFC’s role in encoding expectations and deploying anticipatory signals, we hypothesized that lesions to subregions of PFC would alter CNV sensitivity to predictive context. While both OFC and LPFC are involved in anticipatory processing, their distinct functional contributions renders it challenging to formulate precise hypotheses about how each lesion location would differentially affect the CNV. However, building on our previous findings^104^ and the cognitive map accounts of task space, we expected a superior role for OFC relative to LPFC, evident as a stronger lesion effect on the CNV (i.e., a greater decrement in expectancy-related negativity). In line with proposals that LPFC computes global-level predictions, and the OFC maintains task-state information, we further hypothesized that task-state signals from OFC inform LPFC to deploy global predictions,^110^ consistent with a rostro–caudal feed-forward organization of PFC.^111,112^

## Results

Twenty-two patients with focal lesions (i.e., surgically resected tissue due to tumor in all but two patients who had lesion due to traumatic brain injury) in either the orbitofrontal (OFC, n = 12) or lateral prefrontal cortex (LPFC, n = 10), fifteen age-matched healthy controls (CTR), and seven patients with drug-resistant epilepsy undergoing SEEG recording in the OFC and LPFC (see Material and methods, and ***Figure 1B*** and C), performed an auditory local-global deviance detection task. For every task block, a common tone sequence (presented ∼75%) induced a global expectation (i.e., prediction). On rare occasions, two types of deviant sequences (presented ∼12.5% each) violated the global task rule (i.e., prediction violations). Importantly, only the last tone of each sequence (or its absence) revealed whether a sequence deviated from the global rule. The tone sequences were systematically categorized into five steps (referred to as *expectancy steps*), based on their numbering (varied from 1 to 5) from the last deviant sequence. Participants were instructed to attend to the stream of sounds, count deviant sequences, and report their count in writing at the end of each block (see Material and methods, and ***Figure 1A***). During task performance, we recorded scalp EEG from the prefrontal lesion patients and the CTRs, and SEEG from the patients with epilepsy. An additional group of twenty-two healthy participants (BEH; Behavioral Group) performed a modified version of the experiment (without EEG), where their task was to respond by button press as accurately and fast as possible when deviant sequences were detected. ***Table 1*** presents an overview of the studies included in this report.

**Figure 1.**
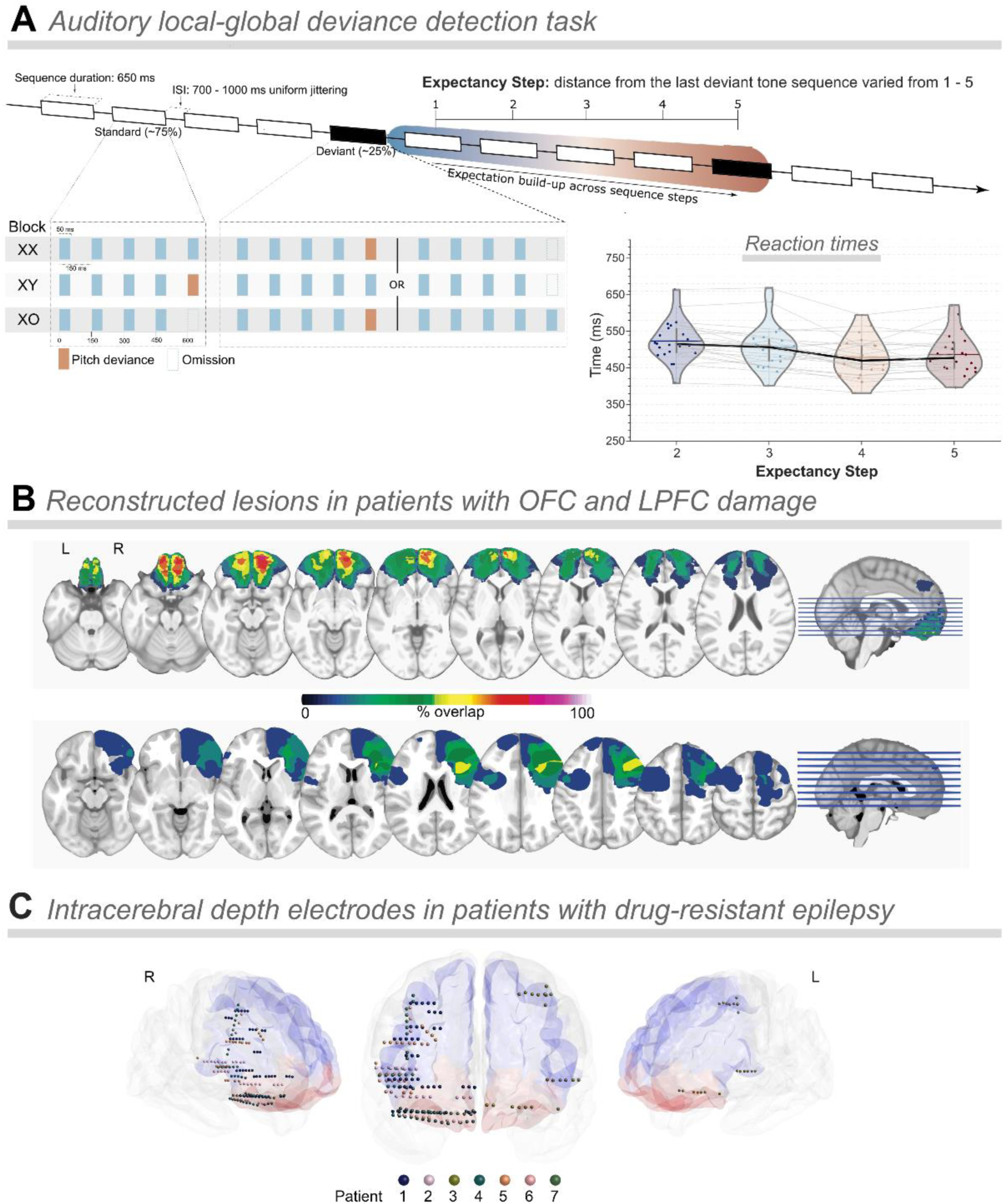
Illustration of task design, lesion reconstruction in patients with frontal damage, and depth electrode coverage in patients with epilepsy. **(A)** *Auditory local-global deviance detection task.* Sequences of 50 ms-duration tones were presented with a fixed stimulus onset asynchrony (SOA) of 150 ms and a uniformly jittered inter-trial interval (ISI) of 700-1000 ms. Each block began with a habituation phase, consisting of 20 identical tone sequences, which would occur commonly throughout the rest of the block. Then, in the remaining part of the block, deviant sequences (25%) were pseudorandomly interspersed among the standards. In regular blocks (XX), standard sequences (75%) consisted of five repetitions of the same tone. They were interspersed with deviant sequences where the fifth sound was either different in frequency type (12.5%) or was shorter (12.5%). In irregular blocks (XY), standard sequences (75%) had a fifth sound differing in frequency type, interspersed with deviant sequences with five repetitions of the same tone (12.5%) or shorter sequences of four identical tones (12.5%). In omission blocks (XO), standard sequences consisted of four repetitions of the same tone. They were mingled with deviant sequences with five repetitions of the same tone (12.5%) or with the fifth sound at a different frequency (12.5%). The *expectancy step* number of each sequence was defined with respect to the numbering since the last deviant sequence. *Reaction times*. Increased speed of global deviant’s detection with expectancy. Violin plots depicting mean reaction times (RT) in milliseconds (ms) across the four *expectancy steps* (2-5) in the RT version of the task conducted with healthy participants (BEH group). Each violin plot represents the distribution of RTs for each step, with individual participant data points connected by gray lines. As the steps increased, a trend towards faster RTs was observed, indicating enhanced response efficiency with increased expectancy. **(B)** *Reconstructed lesions in patients with OFC and LPFC damage.* Lesion overlap for orbitofrontal cortex (OFC: top; n = 12) and lateral prefrontal cortex (LPFC: bottom; n = 10) patients. Aggregate lesion overlay maps per group in axial view. The color code (from 0 to 100%) indicates the percentage of shared lesion coverage across patients. The redder the color, the greater the lesion overlap. Neurological convention was used with the right side of the brain being displayed on the right side of the image and vice versa. **(C)** *Intracerebral depth electrodes in patients with drug-resistant epilepsy*. Normalized space projection of the 202 recorded electrodes onto the 3D glass brain reconstruction of the orbital (red surface; electrodes = 90) and lateral (blue surface; electrodes = 112) PFC. The color-coding of electrodes indicates the individual patients. Note that, by chance, there were more patients with right-than left-hemisphere electrode coverage during the study period (right: n = 6; left: n = 1).

**Table 1.**
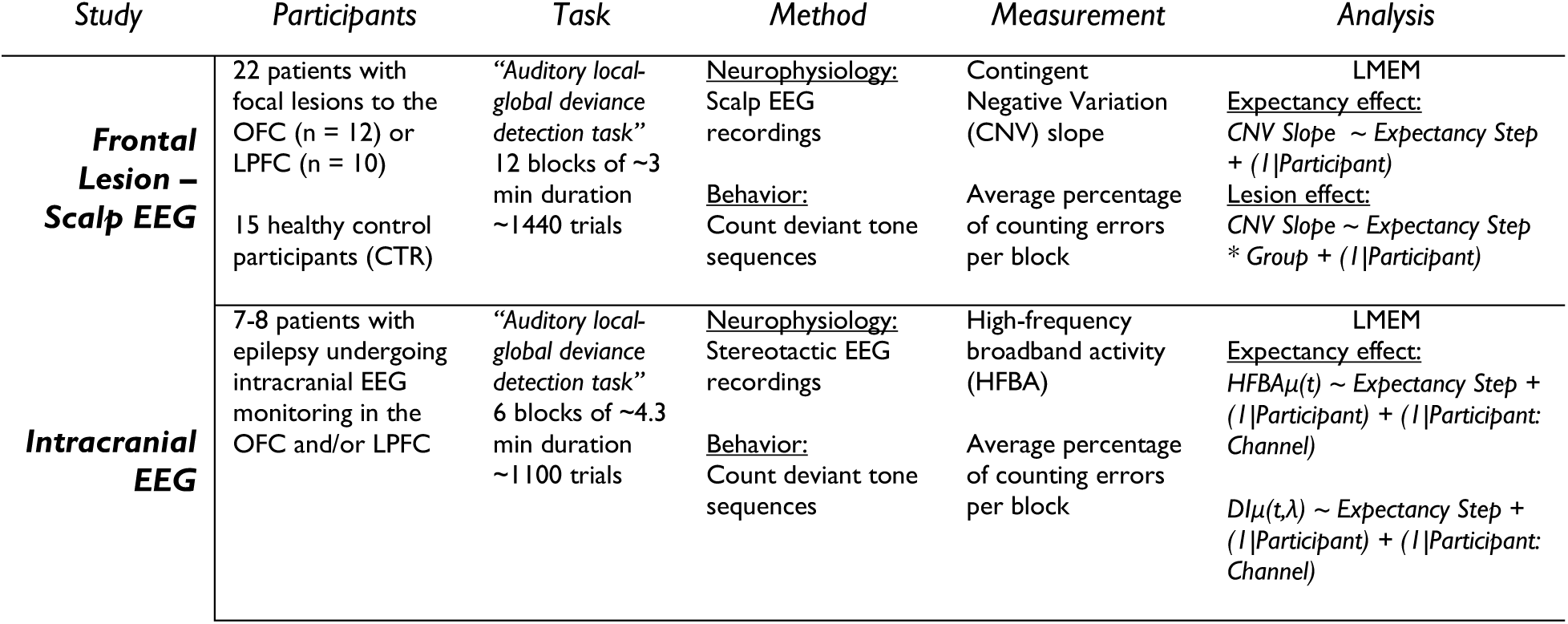
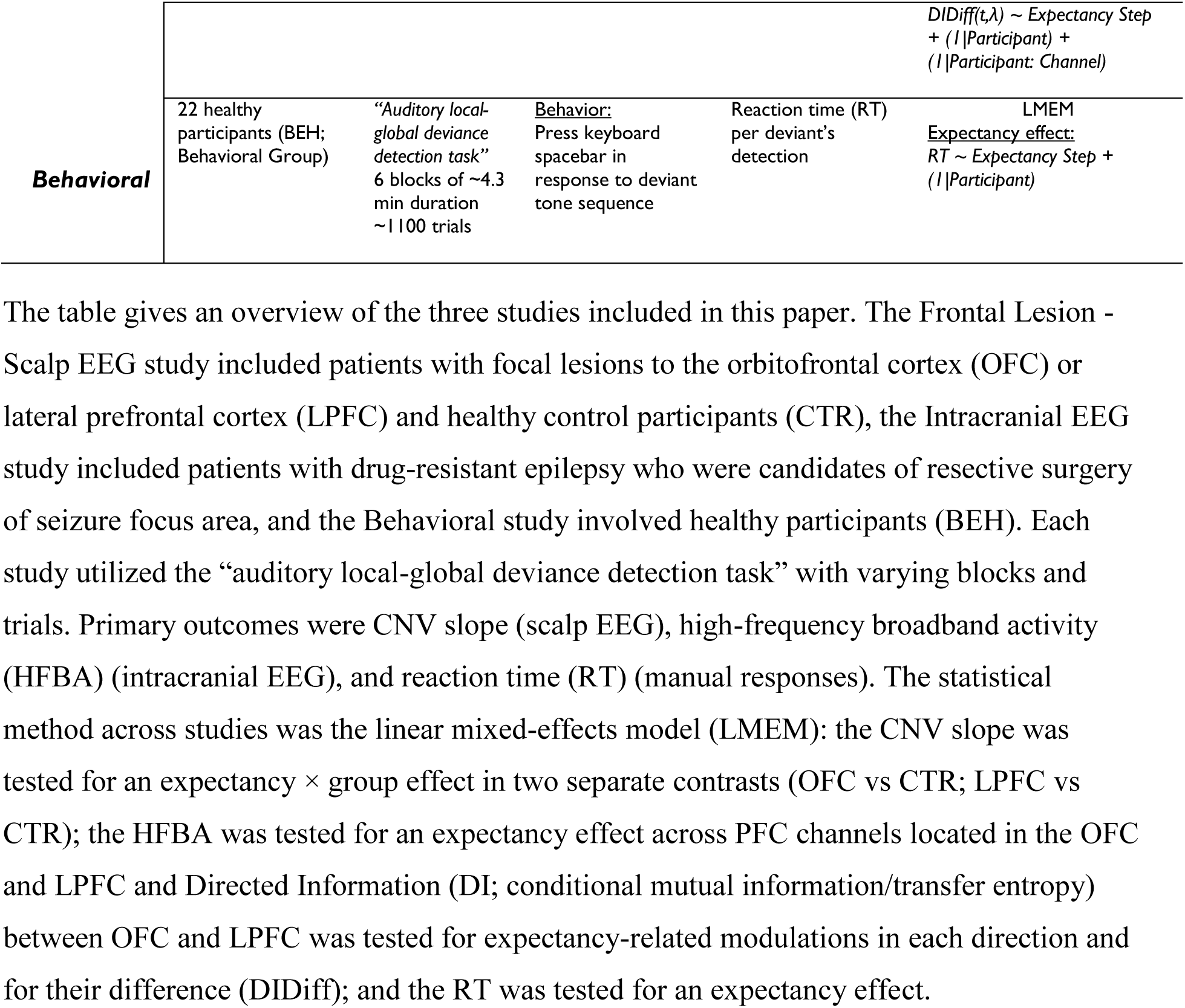
Overview of the studies.

### Expectancy increases detection speed of deviant sequences

The healthy Behavioral Group (BEH) successfully performed the reaction time (RT) version of the task with an error rate of 3.41% (SD = 1.35), and an average RT of 504 ms (SD = 55 ms). Based on previous literature,^113,114^ we expected the RT to be sensitive to the *expectancy step* manipulation. To confirm this hypothesis, a linear mixed-effects model (LMEM) was used with the RT to the detection of global deviants as the dependent variable, the *expectancy step* as the fixed effect predictor, and subject as a random intercept. A significant decrease in RT was observed when the step number increased (*β* = −0.013, *p* < 0.001); *expectancy step 2*: mean RT 524 ms (SD = 55 ms), *expectancy step 3*: 507 ms (SD = 60 ms), *expectancy step 4*: 479 ms (SD = 53 ms), *expectancy step 5*: 487 ms (SD = 56 ms), suggesting that participants’ RTs became faster (until step 4) as the expectancy increased (***Figure 1A***).

### CNV slope encodes the expectancy of global deviants in healthy participants

The number of sequence steps from the previous deviant sound sequence holds predictive information about the outcome of the current sequence (i.e., how probable it is that the current sequence is a deviant). Anticipatory processing before the actual occurrence of a global deviant was quantified by the CNV slope using a least-squares linear fit method^22^ over the sequence interval (0 - 600 ms relative to sequence onset; i.e., the period before the last tone starts). Using LMEM analysis with the *CNV slope* per trial as the dependent variable and the *expectancy step* as the predictor, we first examined whether the slopes of healthy adults differed across trials given the task’s predictive information.

The model (*CNV slope ∼ Expectancy Step + (1 | Participant)*) revealed a robust effect of *expectancy step* in the CTR group across all electrode channels except one after False Discovery Rate (FDR) correction for multiple comparisons. Regression coefficients ranged from *β =* −0.20 at Fpz (*t* = −1.38, *p* = 0.17) to *β =* −0.99 at CP2, (*t* = −6.54, *p* < 0.001), 95% confidence interval (CI). Full per-channel statistics are provided in Supplementary Table 1 and the topographic map of *t*-values, with electrodes surviving the FDR-corrected significance threshold marked in ***Figure 2C***. These results demonstrate a robust effect of the expectancy task information (i.e., *expectancy step*) on the anticipatory neural activity prior to the onset of global deviants. ***Figure 2A*** depicts the CNV waveforms for early (*expectancy step* = 2, in blue) and late (*expectancy step* = 4 and 5, in brown) tone sequences, and the corresponding topographic scalp maps for the three groups of participants.

**Figure 2.**
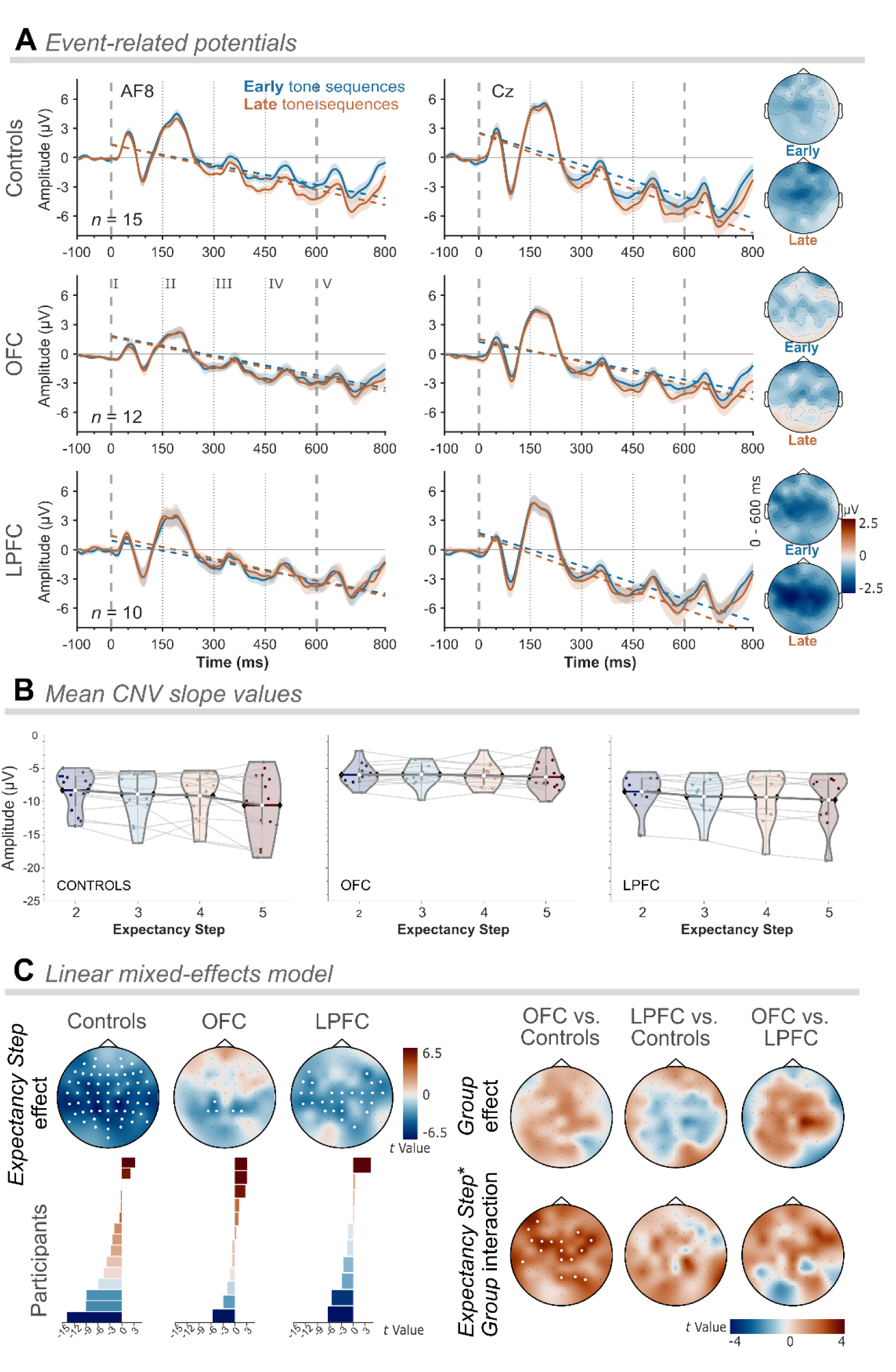
Scalp-recorded contingent negative variation (CNV) was modulated by global deviant expectancy in healthy adults and was significantly reduced in patients with lesions to the orbitofrontal cortex (OFC). **(A)** *Event-related potentials.* On the top, healthy control participants’ (Controls) grand average CNV waveforms at two exemplary electrodes, AF8 (left) and Cz (right). CNV waveforms elicited by early tone sequences (expectancy step = 2) are in blue, and in brown for late tone sequences (expectancy step = 4 and 5). In the middle, orbitofrontal cortex (OFC) lesion patients’ grand average CNV waveforms are shown at the same electrodes and for the same task conditions. At the bottom, lateral prefrontal cortex (LPFC) lesion patients’ grand average CNV waveforms are depicted at the same electrodes for the same conditions. Dashed lines indicate the average linear regression from where CNV slopes were extracted. Vertical lines denote tone onsets. Scalp topographies of voltage averaged in the 600 ms time window from the onset of the first tone to the onset of the fifth tone for early and late tone sequences show a fronto-central negativity maximum, which is more negative for late compared to early tone sequences. **(B)** *Mean CNV slope values.* Violin plots of mean slope value over expectancy steps (2-5) obtained from fronto-central electrodes (i.e., AF7, AF8, Fz, FC3, FCz, FC4, C3, Cz, and C4) for the Controls and the OFC and LPFC lesion groups. Lines connect slope values (points) from the same participant. **(C)** *Linear mixed-effects model.* On the left, scalp topographies show the t-values distribution of the expectancy step effect on the CNV slope for the Controls, the OFC, and the LPFC patients, respectively, when modeled independently. To the right, the scalp topographies illustrate the effect of group (top) and the interaction effect of expectancy step and group (bottom) for OFC vs. Controls (left), LPFC vs. Controls (middle), and OFC vs. LPFC (right). Below the expectancy step effects, participants’ t-values calculated for fronto-central electrodes (see B) obtained with individual linear models.

### Focal lesions to the OFC compromise the expectancy modulation of the CNV slope

We next tested the hypothesis that damage to the OFC would disrupt the CNV response. In contrast to the healthy adults, only five channels showed a significant (FDR-corrected) progressive increase in the CNV slope by *expectancy step*. The model (*CNV slope ∼ Expectancy Step + (1 | Participant)*)’s regression coefficients ranged from *β =* 0.0005 at AFz (*t* = 0.003*, p =* 1) to *β =* −0.60 at CP1 (*t* = −3.55, *p =* 0.025), 95% CI. Full per-channel statistics are provided in Supplementary Table 2 and topographic map of *t*-values, with electrodes surviving the FDR-corrected significance threshold in ***Figure 2C***. To compare the CNV between CTRs and OFC patients, we fitted an LMEM with *CNV slope* as dependent variable and *expectancy step* and *group* (i.e., OFC vs. CTR) as predictors. The model (*CNV slope ∼ Expectancy Step * Group + (1 | Participant)*), where *Group* was treatment-coded with CTR as the reference level, revealed no significant main effect of group, *β* values ranged from *β =* −0.004 at POz (*t* = −0.003, *p* = 1) to *β =* 2.28 at C2 (*t* = 1.7, *p* = 1 (POz), 95% CI. However, it showed a significant interaction between group and *expectancy step* at a cluster of 17 channels after FDR correction, where *β* values ranged from *β* = 0.06 at O1 (*t* = 0.27, *p* = 0.79) to *β* = 0.74 at C4 (*t* = 3.49, *p* < 0.001), 95% CI. Per-channels statistics and *t*-values topographic map are provided in Supplementary Table 4 and ***Figure 2C***, respectively. These results indicate that damage to the OFC impacts CNV modulation in relation to the predictive information of the task, specifically attenuating the normally steeper (i.e., more negative) CNV slope that accompanies higher expectancy.

### Focal lesions to the LPFC moderately decline the expectancy modulation of the CNV slope

We also examined the effect of LPFC damage on the CNV response. When applying the same LMEM with *CNV slope* as the dependent variable and *expectancy step* as the predictor, a significant main effect of expectancy step emerged at 27 electrodes after FDR correction, markedly fewer than in CTRs (63 electrodes), but still showing meaningful modulation. *β* values ranged from *β* = −0.011 at P10 (t = −0.07, *p* = 0.95) to *β* = −0.80 at F6 (t = −3.99, *p* < 0.004), 95% CI, indicating a topographically altered expectancy-driven modulation of the CNV slope relative to the widespread effect seen in healthy adults (Supplementary Table 3; ***Figure 2C***). To quantify this variation between groups, we ran an LMEM with *CNV slope* as dependent variable and *expectancy step* and *group* (i.e., LPFC vs. CTR) as predictors. The model (*CNV slope ∼ Expectancy Step * Group + (1 | Participant)*) showed that the main effect of group was not significant, *β* values ranged from *β* = 0.0008 at CP5, (*t* = 0.001, *p* = 1) to *β =* −2.72 at CP2, (*t* = −1.77, *p* = 0.08), 95% CI. While there was an interaction effect between *group* and *expectancy step* at only six channels (*β* values ranged from *β* = 0.002 at FC4 (*t* = 0.007, *p* = 0.99) to *β* = 0.68 at CP2 (*t* = 2.74, *p* = 0.006), 95% CI), the effect did not survive the FDR correction (Supplementary Table 5; ***Figure 2C***). Where present, the positive interaction *β* indicates that the negative slope (normally produced by higher expectancy) was attenuated in LPFC patients. However, these results preclude strong claims about the difference in how the LPFC and CTR groups integrate the task predictive information to facilitate deviance detection.

Comparing the CNV slopes of the two lesion groups, the model (*CNV slope ∼ Expectancy Step * Group + (1 | Participant)*), where *Group* was treatment-coded with LPFC as the reference level, showed a main effect of *group* at five channels, *β* values ranged from *β =* −0.05 at PO8 (*t* = −0.02, *p* = 0.98) to *β =* 4.90 at CP2 (*t* = 3.32, *p* < 0.001), 95% CI, with no channels passing the FDR correction. The positive *β* at CP2 indicates that OFC patients exhibited a flatter (less negative) CNV slope than LPFC patients. The *Expectancy Step* × *Group* interaction reached significance at nine channels, *β* values ranged from *β =* −0.011 at PO3 (*t* = −0.05, *p* = 0.96 to *β =* 0.75 at F6 (*t* = 2.82, *p* = 0.005), 95% CI, but none of them survived the FDR correction (Supplementary Table 6; ***Figure 2C***). Where present, the positive interaction coefficients imply that the expectancy-driven steepening of the CNV was attenuated in OFC relative to LPFC patients.

### High-frequency broadband activity from OFC and LPFC encodes the expectancy of global deviants in distinct time-windows

To understand the brain areas and neurophysiological processes involved in deviance prediction with higher spatial and temporal resolution, we extracted the high-frequency broadband activity (HFBA; 65 to 145 Hz) directly recorded from OFC (n = 90) and LPFC (n = 112) channels. HFBA is a proxy of the underlying neuronal population within a few mm from the recording electrode.^115–117^ HFBA was extracted for each trial, electrode, and participant (−200 to 800 ms from sequence onset). For each 10-ms time bin, LMEMs were separately fitted to the HFBA (*HFBAμ(t) ∼ Expectancy Step + (1|Participant) + (1|Participant: Channel)*) for the two prefrontal areas, revealing that each area exhibited activity modulated by the *expectancy step* within distinct time windows. ***Figure 3A*** shows the average HFBA signals recorded from OFC (top left plot) and LPFC channels (top right plot), with early tone sequences (*expectancy step* = 2) depicted in blue, and late tone sequences (*expectancy steps* = 4 and 5) shown in brown. The LMEM revealed that the *expectancy step* significantly modulated HFBA in the OFC in an early interval between 180 and 230 ms after sequence onset, with HFBA increasing with increasing *expectancy step* (*p* < 0.05, FDR-corrected). Similarly, the temporal dynamics differed slightly in the LPFC, with significant modulation of HFBA by *expectancy step* occurring later, between 450 and 600 ms post-sequence onset (*p* < 0.05, FDR-corrected). These modulations are indicated by the gray-shaded area in ***Figure 3A***, where the *t*-values from the LMEM exceeded the significance threshold (lower plots). These results show that both the OFC and LPFC exhibited significant HFBA modulation in response to changes in *expectancy step*, with a temporal profile indicating a potentially dynamic interplay between these regions. Specifically, the OFC showed peak sensitivity predominantly during the first half of the tone sequence, while the LPFC was more engaged in the second half, suggesting that these regions reflect distinct time periods during the predictive encoding of the tone sequence.

**Figure 3.**
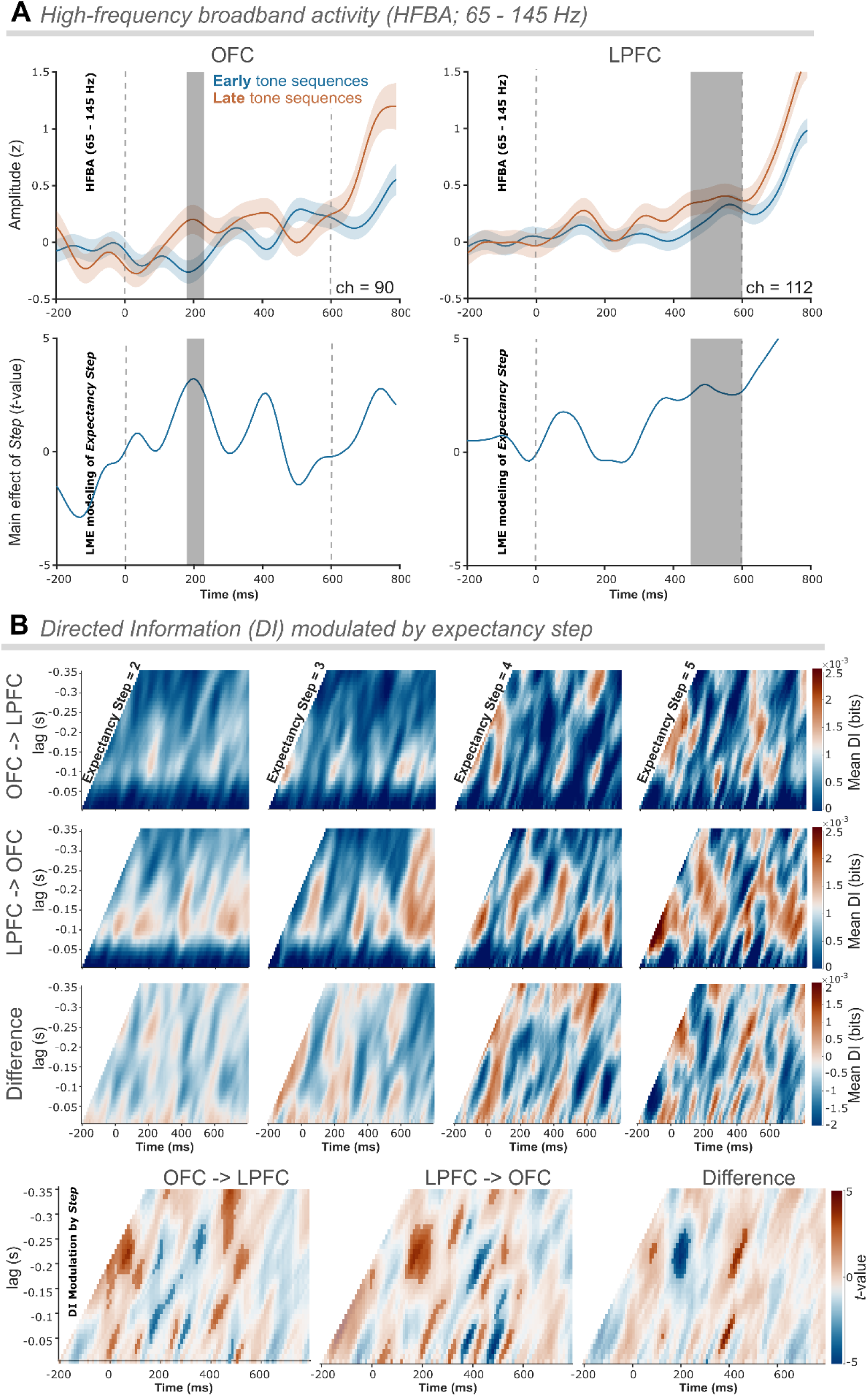
Intracerebral high-frequency broadband activity (HFBA) recorded from orbitofrontal and lateral PFC is modulated by deviance expectancy. **(A)** *High-frequency broadband activity (HFBA; 65 - 145 Hz).* Average HFBA signals recorded from orbitofrontal (OFC; left) channels (n = 90) and lateral prefrontal (LPFC; right) channels (n = 112). HFBA elicited by early tone sequences (*expectancy step* = 2) are in blue, and HFBA signals elicited by late tone sequences (*expectancy step* = 4 and 5) are in brown. Gray area denotes a significant modulation of *expectancy step* in the linear mixed-effects (LME) modeling. **Bottom:** Temporal LME modeling *t*-values time course (−200 - 800 ms) with marked significant segments (gray vertical bar) of *expectancy step* modulation for OFC (left) and LPFC (right). **(B)** *Directed Information (ID) modulated by expectancy step.* The color scale reflects the strength of information flow, with warmer colors indicating higher DI values. **First row:** average DI flow between OFC and LPFC channel pairs (n = 1482); **Second row:** average DI flow between LPFC and OFC channel pairs (n = 1482); **Third row:** average difference (OFC → LPFC minus LPFC → OFC) DI flow. **Bottom row:** LME modeling *t*-values time course (− 200 - 800 ms) showing the modulation of DI flow by *expectancy step* between OFC and LPFC (left), between LPFC and OFC (middle), and the DI difference (OFC → LPFC minus LPFC → OFC; right). Positive DI differences indicate OFC lead (brown), whereas negative values indicate LPFC lead (blue). Significant modulation of the information flow by *expectancy step* (statistically significant *t*-values, uncorrected for left and middle plots, FDR-corrected for right plot) are represented by the opaque colored areas, while non-significant results are shown as transparent areas.

### Expectancy modulates Directed Information (DI) flow between OFC and LPFC asymmetrically

We used Directed Information (DI), which quantifies directional connectivity between neural signals, to assess the dynamic relationship between neural activity in the OFC and LPFC. DI is a statistically robust information theory metric when applied to neural data.^118^ DI quantifies effective connectivity by measuring how well the past activity of a “sender signal” X (e.g., HFBA from OFC channels) predicts the future activity of a “receiver signal” Y (e.g., HFBA from LPFC channels), while accounting for the past activity of the receiver signal Y. Positive values of DI (measured in bits) between signal X at an earlier time and signal Y at a later time indicates that signal X contains predictive information about the future state of signal Y. We computed DI flow for all channel combinations from OFC to LPFC and vice versa within subjects and across the four *expectancy steps* (2-5). Mean DI values are presented in the first two rows of ***Figure 3B***. As a preliminary step, the DI flow between the OFC and LPFC channels was analyzed to examine how the *expectancy step* influences directional connectivity. LMEMs were applied separately to DI information flow in both directions (i.e., from OFC to LPFC and from LPFC to OFC) with *DI* as the dependent variable, *expectancy step* as the predictor or fixed effect and *participant* and *channel* as nested random effects, *HFBAμ(t) ∼ Expectancy Step + (1|Participant) + (1|Participant: Channel)*. ***Figure 3B*** (bottom row) illustrates the results of the LMEMs. The DI flow from OFC to LPFC showed several periods with significant modulation (*p* < 0.05, uncorrected) with the first around 0 to 100 ms after the sequence onset, suggesting that the OFC initially drives expectancy information flow towards the LPFC, indicating that the OFC has a leading role in the process. The LPFC to OFC direction showed several periods with significant modulation (*p* < 0.05, uncorrected), the first around 100-250 ms after the sequence onset, suggesting that LPFC influences OFC later in the sequence processing.

We addressed whether OFC, LPFC, or both lead expectancy-related information flow dynamically. The third row of ***Figure 3B*** depicts the difference in DI flow between the two directions (i.e., OFC → LPFC minus LPFC → OFC), highlighting asymmetry between the two prefrontal areas. Across the *expectancy steps*, varying degrees of positive and negative DI difference values were observed, indicating periods where one direction of information flow was more dominant. To test whether the DI asymmetry was modulated by *expectancy step* in a particular direction,^119^ we fit LMEM with *DI difference* as the dependent variable, *expectancy step* as the fixed effect, and *participant* and *channel* as nested random effects (*DI_Diff_* _(t,λ)_ ∼ *Expectancy Step* + (1|*Participant) +* (1|*Participant: Channel)*). The DI asymmetry modulation showed the dynamic interplay between OFC and LPFC, suggesting a bidirectional interaction between them (***Figure 3B***, bottom row, right plot). The OFC initially led the information flow shortly after the sequence onset (0-100 ms), while at around 200 ms, the LPFC became the dominant source of information flow (*p* < 0.05, FDR-corrected). Finally, the OFC regained dominance between 350 to 420 ms. These shifts in directional dominance suggest a dynamic integration process of expectancy-related information between the two regions.

## Discussion

We asked how OFC and LPFC contribute to the building of implicit expectations that guide predictive and optimal behaviour. Participants performed an auditory local–global deviance detection task in which deviant tone sequences occurred at unpredicted intervals (steps), with the number of standard sequences between them varying, allowing expectancy to increase with elapsed standards. Participants were instructed to detect these global violations, and either count them in the lesion and intracranial studies or respond to them by button press in the follow-up behavioral study, see ***Table 1***. Leveraging previous findings,^22^ we anticipated that deviant expectancy would engage anticipatory neural processing. We analyzed scalp-EEG recorded CNV activity from healthy individuals and patients with focal OFC or LPFC lesions, and intracerebral SEEG activity from the same prefrontal regions in epilepsy patients. In healthy participants, the CNV responses were sensitive to the degree of expectancy of the upcoming deviants. Relative to controls, patients with OFC damage showed a significant attenuation of expectancy-related CNV slope modulation, consistent with a flatter (less negative) CNV at higher expectancy. LPFC patients retained expectancy-related CNV modulation, but it was weaker and more spatially restricted (27 electrodes vs. 63 in controls), and group differences from controls did not survive correction for multiple comparisons. Contrast between the two lesions groups revealed no robust differences after correction, though expectancy effects tended to be flatter in OFC than LPFC. In converging evidence, SEEG-recorded OFC and LPFC HFBA reliably encoded deviancy expectancy, with distinct time windows in the two regions. OFC showed an earlier expectancy-related modulation after sequence onset, whereas LPFC showed a later modulation within the same sequence. Extending this pattern, analyses of directed information flow between OFC and LPFC suggested that expectancy also shaped the direction and timing of inter-regional exchanges. Across the sequence window, interactions appeared bidirectional but asymmetric, with a tendency toward an early OFC→LPFC lead, a mid-interval LPFC→OFC phase, and a later shift back toward OFC→LPFC. To investigate whether expectancy facilitates deviance detection by accelerating manual responses, we conducted a supplementary reaction time (RT) version of the task (BEH group). These results revealed that participants detected deviants faster when they were more expected (i.e., with increasing *expectancy steps*). Taken together, these results provide evidence that prefrontal regions extract implicit environmental regularities to build predictions that tune the brain for optimal detection of changes in the auditory stream.

In the present task, there were no explicit predictive cues; deviant likelihood was a latent state determined by the task’s probabilistic structure, not directly recoverable from the current or immediately preceding stimuli. One possibility is that the neural network exploits the individual sequence-to-sequence transition probability (e.g., using a Markov chain model, successor representation, or state representation)^45,120^ to adjust anticipatory activity towards the auditory input stream. This process requires forming and maintaining an internal abstract representation of the entire pool of states and their transitions (i.e., task context, or “cognitive map”), which reflects the ability to implicitly learn the probability distribution of sequence-to-sequence transitions along the entire experimental block.^53,110^ This anticipatory activity could at the same time be driven by deeply encoded context invariant regularities, such as the increasing probability of state change over time (i.e., a hyper-prior or a fundamental belief).^121,122^ Many events are cyclical and exhibit monotonically increasing probabilities over time, such as changes in traffic signals or arrival times of public transport. For example, the likelihood of a green light rises the longer it has been red and, similarly, the likelihood of a bus arriving increases as time passes. Using a deeply encoded prior represents a computationally efficient solution to regulate anticipation of events, which are cyclically jittered over short time intervals, such as deviant detection in an oddball paradigm. By this scheme, the neural circuits only need to keep track of the time elapsed since the previous deviant sequence. Converging evidence from several studies^45,50,53,54,56,110,120^ suggests that the former hypothesis is the most plausible and will be explored in the following subsections.

### CNV reflects anticipatory activity informed by implicit predictive information

The CNV developed prior to the fifth tone (i.e., the tone that determines deviance) within the sequence and indexed the preparatory state, which facilitates the detection of an upcoming global deviant. Consistent with prior work,^22^ scalp CNV increased with the steps elapsed since the last deviant sequence. Complementing this neural effect, RTs were shorter when deviant were more expected, indicating that expectancy-driven preparation translates into faster detection, in accord with previous studies showing that expectancy speeds RT as event likelihood increases.^123–126^ Together, these neural and behavioral effects indicate that the brain tracks the number of elapsed sequences since the last deviant, in accordance with a global task-state representation,^46,51,127,128^ to predict an upcoming deviant and facilitate its detection. Notably the CNV arose without an explicit cue or request for a motor act and within a brief time window of 600 ms. Unlike classical “S1-S2” CNV paradigms, where a cue (S1) signals a forthcoming imperative stimulus (S2) requiring a motor response, our cue- and response-free design builds expectancy only from latent regularities in stimuli structure. Moreover, while classical S1–S2 CNV paradigms use long intervals that allow the development of the CNV, in this paradigm the CNV develops during a shorter tone sequence, demonstrating that anticipatory activity shaped by implicit predictive information is generated dynamically over short intervals. CNV modulation suggests that the brain tracks increasing expectancy of global deviants across successive standards, reflecting higher-order predictive encoding that primes circuits for detecting and counting task-relevant stimuli, thereby facilitating responses to global violations. Previous studies using auditory local-global paradigms proposed the anterior frontal cortex as a potential driver of this type of CNV,^22,102^ suggesting that the anticipatory process in such tasks might be engaged differently compared to typical sensorimotor CNVs.^71,100,129^ However, scalp EEG source localization is limited in resolving deep generators such as the OFC.

### Converging lesion and intracranial evidence for predictive encoding of deviant tone sequences in the OFC

For the OFC patients, only a limited number of channels showed a significant progressive enhancement of the CNV by deviants’ expectancy, supporting OFC involvement in top-down predictive processing. In complementary intracranial evidence, SEEG-recorded HFBA from OFC channels was significant enhanced by *expectancy step* in an early time window (180– 230 ms post-sequence onset), indicating engagement of the OFC in encoding expectancy-related information during the initial phase of the tone sequence. While these intracerebral signals may reflect processes that also shape the scalp CNV, they arise from different physiological sources and spatial scales (i.e., HFBA indexes local population firing; CNV reflects low-frequency postsynaptic potentials). Accordingly, we interpret them as convergent evidence for expectancy coding rather than a direct equivalence. A direct correspondence would require simultaneous scalp-intracranial EEG recordings.

Although PFC is considered a central hub in the frontoparietal–subcortical network that generates the CNV, no lesion study has directly probed the OFC’s role in CNV generation during auditory predictive processing. A previous “Go/NoGo” S1-S2 study reported no OFC effect on CNV,^130^ but that paradigm does not rely on state-space representations of perceptually unobservable variables or the deployment of abstract information, unlike our task, which relies on indirect information about the occurrence of deviant sequences. By contrast, human fMRI and intracranial EEG (iEEG) studies using non-probabilistic tasks have shown orbitofrontal and mesial prefrontal contribution to preparatory activity, suggesting that OFC/mesial PFC support cognitive processing and decision-making, whereas supplementary motor area (SMA) and primary motor cortex support motor preparation.^85,97–99^ Volume conduction is typically larger for low-frequency activity, precluding a precise localization of the CNV generators using scalp EEG, while bipolar HFBA allows ∼4 mm error in their localization as used in our study.^115^

We previously reported that OFC damage impairs detection of unexpected auditory stimuli – a process requiring integration of information over extended timescales.^104^ Here, we extend those findings by showing OFC engagement in encoding the task’s predictive context using both OFC lesions and intracranial recordings, consistent with a key role of OFC in tracking environmental regularities and structure to optimize perception and ongoing behavior. The implications of these findings are two-fold. Based on prior work indicating that the OFC represent the current context or situation,^45,46,51,53,131^ we propose that the OFC forms and maintains an internal representation (i.e., predictive model or cognitive map) of the task context. This model modulates event preparation, such as deviance detection, in lower sensory areas. In support of this model, other studies show that the OFC provides information about the current state space to signal learning mechanisms in the brain^132–134^ and direct information signals to downstream areas to reshape and update representational models.^135^ Such top-down signals may template the learning of state transition probabilities in other regions, including sensory areas.^135–137^ Additional work implicates the OFC in inferring event likelihood from the probabilistic context of the task,^54,138–140^ underscoring its role in representing and integrating distinct sources of information. Together with our results, these observations support an OFC role in maintaining an internal representation of the task states and the transition probabilities between these states (i.e., the probabilistic context of the task) to inform preparatory processes.

Second, given the role of the OFC in the formation and retrieval of temporal context^41^ and the time perception difficulties observed in OFC patients,^141–143^ our findings are consistent with an impaired evaluation of the passage of time (i.e., since the last deviant sequence). This aligns with the hypothesis that regular target presentation builds a memory of the target-to-target interval, and that such a memory trace might contribute to the generation of a time-based expectancy that elicits a pre-target CNV.^144^ Accordingly, the OFC may be necessary to use temporal information to construct expectancy of a given event. With among the longest temporal integration windows in the brain, the OFC is well-positioned for this computation.^145–147^ Human lesion studies on time perception and production further affirm that an intact OFC is needed to maintain and update temporal information to sustain a stable representational model of task space.^141–143^ OFC damage also diminishes working memory for temporal order (i.e., tracking the relative recency of events)^148^ and contributes to “reality filtering” (i.e., relating past and present).^149^ In our paradigm, the short-term interval between sequences is a perceptually unobservable state keeping track of passing time since the last deviant and allows predictions about when the next deviant will occur, which is likely compromised by OFC damage.

### Intracranial evidence for predictive encoding of deviant tone sequences in the LPFC

The LPFC patient results showed reduced CNV effects, but to a lesser extent than the OFC cohort. When comparing CNV slopes between the healthy controls and LPFC patients, no significant group effect or interaction with *expectancy step* remained after adjusting for multiple comparisons. Visual inspection suggested differences in how LPFC patients and healthy adults encoded implicit predictive information, but these were not sufficiently robust under our statistical criteria. One factor likely contributing is lesion heterogeneity in the LPFC cohort (see Supplementary Figures 2 and 3): in the left hemisphere, lesions varied greatly along the rostral-caudal axis, whereas the right hemisphere showed a greater lesion overlap. Given the bilateral, distributed architecture of the fronto-parietal control network, LPFC functions may be partially compensable via contralesional homotopic cortex and allied control regions. Such functional compensation, e.g., contralesional LPFC recruitment and reallocation within spared fronto-parietal circuits, could attenuate group differences at the scalp, limiting observable CNV effects.^150–152^

By contrast, SEEG-recorded lateral prefrontal activity robustly tracked expectancy: HFBA increased with higher *expectancy steps*, with a later modulation window (450–600 ms post-sequence onset), indicating temporally precise enhanced LPFC engagement closer to the last tone. These intracerebral findings complement the scalp results, suggesting that LPFC contributes to predictive encoding, but in a focal, later stage that can be diluted at the scalp by lesion heterogeneity, compensation and the CNV’s low-frequency distributed generators, while remaining detectable intracranially with higher spatial specificity.

The ventral part of LPFC (i.e., IFG) is implicated in detecting violations in an auditory stream based on prior experience and contextual information. Intracranial EEG studies examining local and global level auditory prediction violations have pointed to the IFG as a source supporting top-down expectation,^25,67,68^ and modulating deviance detection systems in the temporal cortices.^66,153,154^ IFG also mediates attentional reorienting to salient changes in the auditory environment,^155,156^ and enhances change sensitivity.^157,158^ The dorsal part of LPFC (i.e., DLPFC) is involved in detecting salient events and preparing or inhibiting motor responses via attentional control.^159,160^ Monitoring stimulus occurrence, which leads to decreases in RT, also depends on DLPFC integrity.^161–163^ Moreover, right DLPFC supports sustained attention and phasic alertness,^164,165^ while left DLPFC is engaged during the preparation period by regulating and actively maintaining attention according to the demands of the task.^166^ In this context, the LPFC HFBA increase with higher expectancy supports a role of this area in allocating and maintaining the attentional resources and response readiness as deviance becomes more probable. Converging with this account, a simultaneous EEG-fMRI study revealed DLPFC, IFG, and inferior parietal lobule (IPL) activations tracking the contribution of prior probability to CNV modulation.^167^

### Sequential HFBA activity and dynamic predictive information flow between OFC and LPFC

Expectancy-induced HFBA modulation began earlier in OFC (180–230 ms) than in LPFC (450–600 ms), suggesting a sequential involvement of these regions. We computed directed information (DI) asymmetry, a measure of effective connectivity, to assess the leading direction of information flow between the two regions.^118,119^ Bidirectional flow was modulated by *expectancy step* and unfolded over distinct time windows. OFC led early after tone sequence onset (0–100 ms), directing expectancy-related information to the LPFC. LPFC exerted greater influence later in the sequence (100–250 ms), becoming the dominant source of information flow. OFC then regained dominance between 350-420 ms, indicating a continuous exchange of expectancy-related information. These patterns are consistent with an early OFC influence initiating predictive encoding of auditory sequences and a later LPFC engagement likely supporting task performance (i.e., enhancing predictions about upcoming sequence deviations).

This dynamic interplay between the OFC and LPFC underscores complementary roles with alternating periods of directional dominance. The OFC may form and maintain predictive information via a representational model or “cognitive map,” providing the temporal context needed to initiate expectancy-related processes.^34,46,49,54,128,168^ In turn, the LPFC may allocate attentional resources and maintain phasic alertness to the impending stimulus,^157,158,163,164,166^ leveraging predictive information received from the OFC. This aligns with the proposed rostro-caudal hierarchical model of PFC functioning, in which more rostral portions exert greater influence on more caudal areas in support of progressively more abstract types of action control.^111,112,169^ We further speculate that context encoding by the OFC influences behavior via a cascade of activity along the lateral surface of the PFC.^56,170^ The time-varying information exchange between the two areas stresses the importance of temporal dynamics in shaping the brain’s predictive and inferential processing in audition.

### Limitations and future directions

We acknowledge the low number of patients with intracranial electrodes studied, which is a common constraint in the iEEG literature.^171–173^ This limitation led to only a partial overlap between the regions affected by the PFC lesions and those where electrodes were implanted in patients with epilepsy. Therefore, the lesion and SEEG results might represent divergent contributions from different prefrontal sub-regions. Future research should focus on methods such as effective connectivity and multivariate pattern analyses to better define how the OFC and LPFC communicate during tasks requiring auditory prediction and deviance detection.

### Conclusion

Our study provides evidence that both OFC and LPFC support anticipatory processing of auditory deviant sequences, informed by implicit predictive information. Both OFC and LPFC lesions decreased scalp-recorded CNV activity, with robust effects only observed in the OFC cohort. Intracranial recordings provided additional support for the role of OFC and LPFC in building up expectancy-related activity with high spatiotemporal precision. Modulation of the HFBA recorded from electrodes in those regions tracked the expectancy of deviant tone sequences, with electrodes in OFC showing earlier activity modulation than the ones in LPFC. Based on the latency differences and information flow patterns between the two areas, we speculate that the OFC provides current state information (i.e., predictive task context information) to the LPFC. In turn, the LPFC exerts top-down modulation of sensory processing to facilitate anticipation of deviances within the predictive context, with bidirectional information flow between these regions.

## Materials and Methods

### Stimuli and experimental design

Healthy control participants and patients with frontal brain lesions completed a variant of the auditory local-global paradigm, the *auditory local-global deviance detection task*, which consisted of 12 blocks of stimuli (6 unique block types repeated twice), with breaks after every block. In each block of ∼ 3 minutes duration, five or four tones lasting 50 ms and spaced 100 ms apart were presented in grouped sequences (i.e., trials) lasting 650 ms. The inter-sequence interval (ISI) was randomly jittered from a uniform distribution between 700 and 1000 ms. Individual tones were composed of three sinusoidal tones, either tone type A (440, 880, and 1760 Hz) or tone type B (622, 1244, and 2488 Hz). Within each sequence, the first four tones were always identical, either A or B. The fifth tone was either the same as the preceding ones (i.e., AAAAA or BBBBB), different in pitch (i.e., AAAAB or BBBBA), or omitted (i.e., AAAA_ or BBBB_). Each experimental block began with a habituation phase, consisting of 20 identical tone sequences to establish the global rule, which is the pattern that participants implicitly learn to expect across sequences. In the test phase (∼100 sequences per block), 75% of the sequences continued the global rule and were considered global standards, while the remaining 25% violated it and were classified as global deviants. The local rule, by contrast, is defined within a single sequence: the final tone either follows or violates the repeating pattern established by the first four tones. For example, in a sequence like AAAAB, the final tone violates the local repetition pattern and is considered a local deviant, whereas AAAAA is a local standard. Across all blocks, the total number of tone sequences presented was approximately 1440.

Part of the results reported in the current paper are a reanalysis of data from a previously published study^104^ in which we examined effects of OFC and LPFC lesions on auditory evoked potentials to violation of auditory prediction at two hierarchical levels of task abstraction. In the present report, we extended this analysis by including SEEG patients performing the auditory local-global deviance detection task adapted from the scalp-EEG recording protocol for optimal bedside data acquisition in the hospital environment. To streamline the procedure, block repetitions were eliminated, reducing the number of blocks from 12 to 6. To offset the reduced number of trials, each block included ∼ 60 additional tone sequences. Moreover, the ISI was shortened by an average of 100 ms by negatively shifting the uniform ISI sampling distribution. Conjointly, these changes extended the block duration from ∼ 3 minutes to ∼ 4.3 minutes and decreased the overall run time by ∼ 10 minutes to facilitate better patient performance. Task instructions, stimulus composition, and sequence probabilities corresponded with the scalp-EEG implementation. The total number of tone sequences was approximately 1100.

An additional group of healthy participants performed a modified reaction time (RT) version of the task, where participants were instructed to respond (by pressing the spacebar) as accurately and fast as possible when deviant sequences were detected. This version’s design and length was the same as the one run for the patients with epilepsy, except that instead of counting global deviants, we measured the speed of global deviants’ detection as a behavioral index to infer facilitation of detection with expectancy.

Three block types were defined for all the versions of the task: Regular, Irregular, and Omission. In regular blocks, standard sequences consisted of five repetitions of the same tone and were interspersed with deviant sequences where the fifth sound was either different in frequency type (∼12.5%), or was omitted (∼12.5%). The irregular blocks were similar, except that the standard sequences had a fifth sound differing in frequency type, interspersed with deviant sequences with five repetitions of the same tone (∼12.5%), or shorter sequences (∼12.5%). In omission blocks, standard sequences consisted of four repetitions of the same tone and were interspersed with deviant sequences with five repetitions of the same tone (∼12.5%) or with the fifth sound being different in frequency type (∼12.5%). Deviant sequences were interspersed among standard sequences in a pseudorandom order with one to four standards presented between two deviants ***Figure 1A***.

For the present analysis, the tone sequences were defined with respect to the time elapsed since the last deviant sequence. Specifically, *expectancy step* = 2 occurred after the presentation of a deviant and a standard sequence. *Expectancy step* = 3 occurred after a deviant and two standards. *Expectancy step* = 4 occurred after a deviant and three standards. Finally, *expectancy step* = 5 occurred after a deviant and four standards. The *a priori* distribution of these deviant sequences was approximately 25%, 30%, 10%, and 35% for the second, third, fourth, and fifth position after a deviant, that is, the probabilities of having deviant-to-deviant chunks of length two, three, four, and five steps. In consequence, the conditional (i.e., hazard) probabilities that the next sequence is deviant, given that no deviant has yet occurred, were approximately 25%, 40%, 22%, and 100% for *expectancy step* 2, 3, 4, and 5, respectively. For example, after listening to the second standard sequence, the conditional probability of a deviant to occur (*expectancy step* = 3) is 40%, which is different from the *a priori* probability of 30% for chunks of length 3. The *expectancy step* indexes elapsed standards and the intended rise in expectancy; it serves as an ordinal proxy rather than a numeric probability, given the task’s objective conditional probabilities (25%, 40%, 22%, and 100%). For intuition, a simple monotonic step index, which is a simple linear rescaling showing that expectancy rises with each additional standard, would place step 3 at 50%. See Supplementary Table 9 for an overview of the *expectancy step* indices arising from the task design. For illustrative purposes, we designated the *expectancy step* 2 as *“early”* and the *expectancy step* 4 and 5 as *“late”* (***Figure 1A***).

### Reaction time version of the auditory local-global deviance detection task

The behavioral RT version of the task was introduced later in the project as a supplemental measure of global deviance detection. Given that the contingent negative variation (CNV) was found to track the probability (expectancy) of deviant sequences in an otherwise regular auditory stream, we aimed to elucidate whether the facilitation of deviant detection (reflected in shortened RTs) is modulated by the same underlying probabilistic structure observed in the CNV.

#### Participants

A sample of 27 healthy adult volunteers were recruited for the additional RT study. Two participants were excluded because of age and epilepsy diagnosis, respectively. Three participants were additionally excluded due to being outliers with respect to their RT performance, defined as three scaled median absolute deviation (MAD) away from the median. This resulted in a sample of 22 participants (BEH; Behavioral Group: 12 females; mean age ± SD, 26.3 ± 3.2 years; education: one participant with 11-13 years, 12 participants with 14-16 years, and nine participants with 17+ years; handedness: 18 right-handed, 2 left-handed, and 2 ambidextrous), who were included for the analysis of the task, which required a motor response to deviant tone sequences. All participants reported normal hearing, no neurological problems, and no cognitive difficulties. They also reported not receiving any psychiatric treatment, including no medication for mental illness. Participants gave written informed consent before participation. The study was approved by the Department of Psychology’s internal research ethics committee (University of Oslo), and was conducted in agreement with the Declaration of Helsinki.

#### Reaction Time data acquisition and analysis

To behaviorally measure facilitation of global deviant detection based on expectancy manipulation, we instructed participants to use the index finger of their dominant hand to press the keyboard spacebar each time they identified an uncommon or rare tone sequence. First, RT outlier trials were identified and removed (0.97%) using MATLAB’s *isoutlier* function with the “median” method, which identifies elements more than three scaled MAD from the median, effectively addressing data that may be skewed or non-normally distributed. Then, the RT response for every deviant trial (i.e., tone sequence) and each participant was modeled as a function of expectancy (*Expectancy Step* = 2, 3, 4, 5). We performed linear mixed-effects model (LMEM) analysis (*RT* ∼ *Expectancy Step* + (1|*Participant*)) with *RT* per trial as the dependent variable, the *Expectancy Step* as the predictor or fixed effect, and *Participant* as the random effect. The analysis was conducted in MATLAB using the *fitlme* function.

### Counting performance in the auditory local-global deviance detection task

To maintain participants’ selective attention on the auditory stream and obtain a behavioral measure of deviance detection, we instructed participants to count and report at the end of the block the number of infrequent tone sequences (i.e., deviant sequences) detected per block. We then computed the average percentage of errors over blocks, which is the deviation (positive or negative) relative to the actual number of presented global deviants.

### Scalp electroencephalography (EEG) protocol

#### Patients with acquired frontal brain lesions and healthy controls

Twelve patients with bilateral lesions to the orbitofrontal cortex (OFC: 8 females; mean age ± SD, 47.9 ± 11.7 years; mean education ± SD, 15.0 ± 3.1 years) and ten patients with unilateral lesions to the lateral prefrontal cortex (LPFC: 6 females; mean age ± SD, 40.9 ± 11.8 years; mean education ± SD, 15.9 ± 2.4 years) were assessed for this study. Most lesions resulted from resection surgery; among the traumatic brain injury (TBI) cases, one did not undergo resection, whereas another underwent evacuation of a hematoma with removal of contused tissue. All patients were in the chronic phase of recovery, that is, at least 2 years after injury or surgery. Details about lesions’ etiology, size, and location are provided in **Table *2***. The patients were compared to fifteen healthy control participants (CTR; 10 females; mean age ± SD, 48.5 ± 10.1 years; mean education ± SD, 16.5 ± 2.2 years; no history of neurological or psychiatric conditions). The three study groups did not differ significantly regarding sex, age, or years of education. Moreover, the OFC and the LPFC group did not differ from controls on any of the standardized and normed neuropsychological measures (including estimated IQ, psychomotor speed, cognitive flexibility, and inhibitory control). Group means and statistical comparisons on neuropsychological test measures are reported in Supplementary Table 7.

**Table 2.**
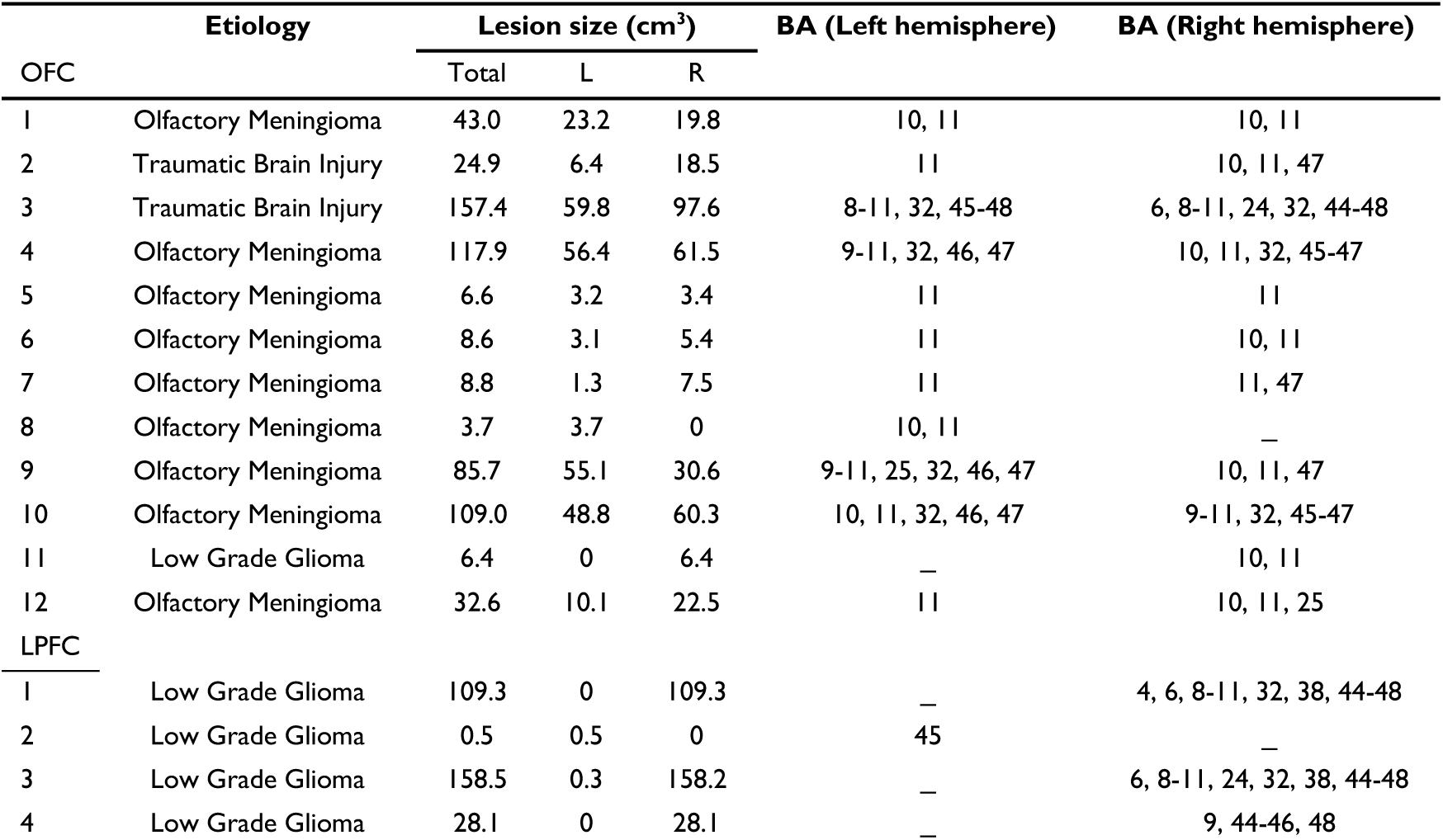
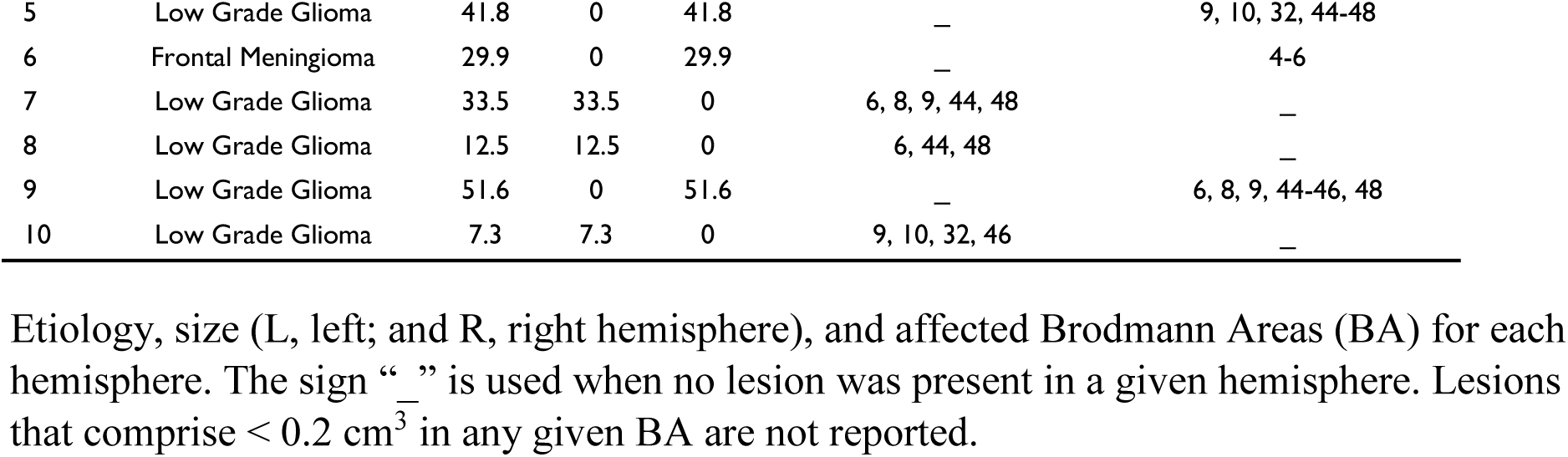
Characteristics of lesions to the orbitofrontal (OFC) and lateral prefrontal (LPFC) cortex.

Healthy control participants received 400 NOK (approximately 50 USD) for participation in the entire research project (neuropsychological assessment, EEG recording, and MRI scanning). Lesion patients participated in conjunction with clinical follow-ups at the hospital’s outpatient clinic. Their travel and accommodation expenses were covered. All participants gave written informed consent before inclusion in the study. The study was authorized by the Regional Committees for Medical and Health Research Ethics (REC), South-East Norway 2014/381 and was conducted in accordance with the principles stated in the 1964 Declaration of Helsinki and its amendments.

#### Lesion mapping

Lesion location and extent were assessed based on structural MRI scans obtained around the time of participation in the study. The lesions were manually drawn on Fluid Attenuated Inversion Recovery (FLAIR) sequence images (1 × 1 × 1 mm^3^ resolution) for each patient’s brain utilizing MRIcron2 (www.mccauslandcenter.sc.edu/mricro/mricron/). High-resolution T1-weighted images aided in determining the boundaries of the lesions, which were subsequently verified by the neurologist and the neurosurgeon in our team (R. T. K and T. R. M). Every participant’s brain was extracted from the T1 image employing the FSL Bet algorithm (FSL3). Thereafter, the brain images were spatially normalized to the Montreal Neurological Institute MNI-152 template space using the unified segmentation and normalization procedures of the Statistical Parametric Mapping software (SPM12: www.fil.ion.ucl.ac.uk/spm/). The outlined lesions were included as masks. Additionally, the transformation matrix was applied to the individual patient’s FLAIR and lesion mask images. The cumulative lesion reconstructions for the OFC and LPFC groups are illustrated in ***Figure 1B***, while ***Table 2*** gives detailed lesion information. Individual lesion maps for each patient are presented in the Supplementary Figures 1, 2, and 3 to show the individual lesion distributions.

#### Scalp EEG data acquisition and pre-processing

We recorded participants’ scalp EEG using a 64-channel Active Two system (BioSemi, Amsterdam, Netherlands). The FieldTrip toolbox^146^ for MATLAB (R2018a, MathWorks Inc., Natick, MA, USA) was used for offline EEG data processing. The continuous EEG data underwent high-pass filtering back and forward (zero-phase) with an infinite impulse-response (IIR) Butterworth filter (order: 3), half-amplitude cutoff at 0.01 Hz. To ensure data quality, we visually inspected the continuous data and identified any channels or segments contaminated with noise, saving this information for later rejection of overlapping epochs. We then employed independent component analysis (ICA) to identify and remove blinks and horizontal eye movement components. Noisy channels were rejected and interpolated from the neighboring electrodes using spherical spline interpolation.^147^ We segmented the pre-processed EEG data into epochs spanning from −1100 ms to 900 ms in relation to the onset of the last tone in the sequence for the Regular and Irregular blocks, or its omission in the Omission block. The epochs generated were down-sampled to 512 Hz and baseline corrected over the 300 ms preceding the onset of the first tone in the sequence.

#### Scalp EEG data analysis

For every electrode and trial (i.e., tone sequence), we computed the slope of the least-squares linear fit over the −600 to 0 ms interval in each subject-wise ERP waveform.^22^ Outlier slope values that were more than three scaled MAD from the median were rejected (CTR: 1.14%; OFC: 1.25%; LPFC: 1.48%). To evaluate the build-up of the CNV slope in individual trials as a function of expectancy (*Expectancy Step* = 2, 3, 4, or 5) across participants in each group (CTR, OFC, and LPFC), we performed LMEM analysis (*CNV Slope* ∼ *Expectancy Step* + (1|*Participant*)), separately for each group. The analysis was conducted in MATLAB using the *fitlme* function. The model incorporated CNV *Slope* per trial as the dependent variable and *Expectancy Step* as the predictor or fixed effect. To account for shared variance in electrodes within a participant, *Participant* was included as the random effect in the model. To account for the multiple comparisons problem, we applied the Benjamini-Hochberg false discovery rate (FDR) method^174^ with α = 0.05 to the results obtained from the model across all channels to correct for multiple comparisons.

To evaluate the effect of OFC or LPFC lesions in the CNV slope modulated by expectancy, we modeled the CNV slope as a function of expectancy (*Expectancy Step* = 2, 3, 4, 5) and group. We performed LMEM analysis (*CNV Slope* ∼ *Expectancy Step* * *Group* + (1|*Participant*)), separately for each group comparison (OFC vs. CTR or LPFC vs. CTR). The model included CNV *Slope* per trial as the dependent variable, *Expectancy Step* and *Group* as the predictors or fixed effects, and *Participant* as the random effect. Although CNV effects are typically fronto-central,^22^ we analyzed all 64 electrode channels and *p*-values from the models were FDR-corrected at α = 0.05. **Figure *2C*** illustrates all the scalp-wide results.

### Stereoelectroencephalography (SEEG) protocol

#### Patients with epilepsy undergoing SEEG monitoring

Seven patients (2 females; mean age ± SD, 42.1 ± 8.1 years; mean education ± SD, 15 ± 2.4 years) with drug-resistant epilepsy undergoing stereotactic electroencephalography (SEEG) for pre-surgical evaluation at Oslo University Hospital were recruited. The inclusion criterion for this study was the presence of electrodes in the orbitofrontal or the lateral prefrontal cortex. All patients had an estimated IQ score above 80. Detailed participant demographics and clinical characteristics are provided in Supplementary Table 8. The study was authorized by the Regional committees for medical and health research ethics - Region North Norway (2015/175) and was conducted in accordance with the principles stated in the 1964 Declaration of Helsinki and its amendments.

#### SEEG data acquisition

The SEEG data were obtained from depth arrays implanted in the lateral and orbital PFC (8-18 contacts/arrays; 3.5 mm inter-electrode distance; DIXI Medical, France) using either an ATLAS (Neuralynx, USA; n = 4) or Brain Quick (Micromed, Italy; n = 3) system with a sampling rate at 16 000 or 2048 Hz, respectively. Field potentials were recorded relative to an adjacent white-matter contact or the average of a set of frontopolar subdermal contacts. To facilitate electrode localization and anatomical labeling, brain images were obtained from each patient pre- and post-implantation, by magnetic resonance imaging (MRI) and computerized tomography (CT), respectively.

#### SEEG electrode localization

The iElectrodes toolbox^175^ was used to acquire the electrode coordinates and their associated anatomical labels based on the co-registered structural images and a parcellation of the cortical surface derived from the Destrieux atlas.^163^ The post-implantation CT image was co-registered to the pre-implantation T1 image based on MRI using SPM12. Electrode coordinates were extracted from the CT images using clustering or the GridFit algorithm.^175,176^ The cortical parcellation was derived by processing the T1 image through the FreeSurfer recon-all pipeline.^177^ Spatial normalization to the MNI-152 template was done in SPM12.^178^

#### Region of interest definition for SEEG electrodes

Lateral prefrontal cortex (LPFC) was defined as the inferior, middle, and superior frontal gyrus anterior to the precentral gyrus, excluding the medial aspect of the superior frontal gyrus. Orbitofrontal cortex (OFC) was defined as the frontal cortex ventral to the callosal genu, including the orbital operculum and excluding the pars orbitalis. The electrode’s anatomical labeling was performed in native space based on probabilistic labels from the Destrieux atlas^179^ and visual confirmation via inspection of the T1-weighted MRI images. In total, there were 112 LPFC electrodes, with an average of 16 electrodes per patient (range: 6 – 26), and 90 OFC electrodes, with an average of 13 per patient (range: 3 – 25).

#### SEEG data pre-processing

Monopolar SEEG traces were visually inspected to assess data quality and to identify epileptiform activity and artifactual signals, which were excluded from subsequent analysis by either removing the affected contact or by discarding samples (resulting in the removal of an average of 12.1%, SD = 14.6%, of the trials across participants). Power line noise was ameliorated by interpolating the spectrum in narrow bands centered on 50 Hz and the first four harmonics.^148^ To achieve high spatial sensitivity, bipolar channels were created by computing the first spatial derivative of pair-wise contacts, moving out from the innermost contact in steps of 1.^115^ Finally, the data were segmented into 3.4 s epochs around the onset of a tone sequence and demeaned epoch-wise.

High-frequency broadband activity (HFBA) was extracted by band-pass filtering each trial in eight non-overlapping bands from 65 to 145 Hz with 10 Hz bandwidth, followed by squaring of the magnitude of the Hilbertized signals. The signals were subsequently down-sampled to 500 Hz. We applied a z-scoring procedure prior to averaging over bands to correct for the 1/f drop-off and streamline subsequent group-level analysis. The eight band traces were normalized using a bootstrapped baseline distribution, which was generated by randomly resampling baseline values 1000 times (from −0.3 to 0 sec relative to sequence onset). The traces were normalized by subtracting the mean of the distribution and then dividing by the standard deviation of the bootstrap distribution.^180^ HFBA signals were smoothed using a 50 ms moving average window and down-sampled to 100 Hz for statistical analysis.^168^ An outlier rejection procedure was implemented to exclude trials that deviated by 7 median absolute deviations from the median for 10 or more consecutive samples (i.e., 100 ms).^181^

#### Modeling HFBA modulation by Expectancy Step

To assess whether neural activity in the LPFC and OFC encodes *expectancy step* predictively, we fitted separate LMEMs for the two areas to the mean HFBA (*HFBA_μ(t)_*) between sequence onset and offset (0 – 600 ms). Since LMEMs effectively account for dependencies in the regressor variables,^182^ the HFBA from all channels and participants in the regions of interest were included in the models. LMEMs were defined with *HFBA* as dependent variable, *Expectancy Step* as predictor or fixed effect, and *Participant* and *Channel* as nested random effects: *HFBA*_μ(t)_ ∼ *Expectancy Step* + (1|*Participant) +* (1|*Participant: Channel)*. The data were transformed using a rank-mapping from the empirical distribution function to the normal distribution function.^183,184^ *p*-values were corrected for multiple comparisons across time using the FDR method with α = 0.05.^174^ Model residuals were inspected to assess conformity of gaussianity using quantile-quantile plots and histograms.

#### Directed Information (DI) analysis between OFC and LPFC

To analyze the causal relationship between the neural activity in the OFC and LPFC we computed the Directed Information (DI, also known as conditional mutual information or transfer entropy) using the GCMI toolbox (Gaussian Copula Mutual Information^118^). DI from X to Y (I (X_t1_; Y_t2_ | Y_t1_)) measures the time lagged dependence between signal X at time t1 and Y at a later time t2, conditioned to the past dependence of the signal Y itself at time t1. DI is a model-free measure that can be interpreted as the information theory analog of Granger Causality.^185^ DI was computed in both directions for all channel pairs between OFC and LPFC for each *Expectancy Step* for the period between 0 and 600 ms after the onset of the tone sequence, and for 0 to 350 ms lags. To avoid interference from responses to previous sequences, the analysis was limited to lag samples up to 200 ms before the sequence onset. Note that no deviant responses were present in the baseline period, as our analysis excluded data from *Expectancy Step = 1*.

LMEMs were used to assess, at the group level, the effect of *Expectancy Step* over the DI measures. As a preliminary analysis, OFC to LPFC and LPFC to OFC modulations were assessed independently. LMEMs were defined with *DI* as dependent variable, *Expectancy Step* as predictor or fixed effect, and *Participant* and *Channel* as nested random effects: *DI*_μ(t,λ)_ ∼ *Expectancy Step* + (1|*Participant) +* (1|*Participant: Channel),* where *t* is the time of interest and λ the time lag.

For the main analysis, we evaluated the periods of significant unidirectional DI modulation to establish whether OFC or LPFC had a leading role (DI asymmetry). The difference *DI_Diff_* between them (i.e., OFC → LPFC minus LPFC → OFC) was computed.^113^ LMEMs were defined with *DI_Diff (t,λ)_* as dependent variable, *Expectancy Step* as predictor or fixed effect, and *Participant* and *Channel* as nested random effects: *DI_Diff_* _(t,λ)_ ∼ *Expectancy Step* + (1|*Participant) +* (1|*Participant: Channel).* The data were transformed using a rank-mapping to ensure normality in the distributions.^183,184^ The *p*-values were corrected for multiple comparisons across time and lags using the FDR method with α = 0.05.^174^

## Acknowledgements

We are very thankful to the patients and healthy participants who participated in this study. We acknowledge Dr. Maja Dyhre Foldal for her contribution to data collection. Professor Per Kristian Hol is acknowledged for his valuable help with the clinical evaluation of the MRI scans of lesion patients and healthy controls. We also thank the EEG technicians at the Department of Neurosurgery, Oslo University Hospital - Rikshospitalet.

## Funding information

Research Council of Norway grant 240389 (AKS, TE, TRM)

Research Council of Norway grant 314925 (AOB, VV)

Research Council of Norway grant 262762 RITMO (AKS, TE)

Research Council of Norway grant 274996 RITPART (AKS, TE)

National Institute of Neurological Disorder and Stroke grant NINDS NS21135 (RTK)

National Institute of Mental Health Conte Center P50 MH109429 (RTK)

## Author contributions

Conceptualization: OA, AKS, AOB

Methodology: OA, VV, AKS, AOB

Formal analysis: OA, VV, SLL, AKS, AOB

Investigation: OA, VV, SLL, IF, AL, JI, TRM, LE, PGL, RTK, TE, AKS, AOB

Resources: PGL, TE, AKS, AOB

Data curation: OA, VV, IF, TRM, PGL, TE, AKS, AOB

Project administration: OA, AKS, AOB

Funding acquisition: TRM, PGL, RTK, TE, AKS, AOB

Visualization: OA, VV, AOB

Supervision: RTK, TE, AKS, AOB

Writing—original draft: OA, AKS, AOB

Writing—review and editing: OA, VV, SLL, IF, AL, JI, TRM, LE, PGL, RTK, TE, AKS, AOB

## Data and materials availability

Materials for the Experimental scripts and task stimuli, custom analysis code, and a processed version of the datasets are available at https://osf.io/su3pg/

The conditions of the ethical approval of this study do not permit public archiving of de-identified raw data. Neither the patients nor the healthy control participants have consented to making their data publicly available. The reason for the restrictions concerning public sharing of clinical data is that the patient samples are small given the relative rareness of individuals with focal brain lesions or SEEG recordings, and thus the constellation of demographic and clinical information results in an increased risk of patients being identified.

Interested researchers seeking access to the original de-identified data supporting the claims in this paper would have to submit a short study plan of the proposed research to the PI of the project and lesion registry, Anne-Kristin Solbakk. The study plan would be evaluated by the project PI, the head of the Department of Neurosurgery at Oslo University Hospital, and the head of research at the Department of Psychology. Next, the PI would ask REC for permission to share de-identified data with the researcher/institution. After REC approval, the head of research at the Department of Psychology, the Data Protection Officer at Oslo University Hospital, and the other interested party would sign data transfer agreements, before data transfer would take place. The data can be accessed and used only for academic purposes.

## Competing interests

The authors declare no competing interests.

## Notes

### Competing Interest Statement

The authors have declared no competing interest.

