## Supplementary Material for "Predictive encoding of auditory sequences in the human prefrontal cortex"

### SUPPLEMENTARY INFORMATION

### FRONTAL LESION STUDY

**SCALP RECORDED CNV SLOPE:** Linear mixed-effects model results across conditions and participant groups.

The following tables summarize the linear mixed-effects (LME) model results for each of the 64 EEG electrodes in the CTR, OFC, and LPFC group. The model tested the effect of the *expectancy step* (2-5) on trial-wise CNV slope. The tables report the regression coefficient (column name: *b*Value), *t*-statistic (*t*Stat), uncorrected *p*-value (*p*Value), and FDR-adjusted *p*-value (adjP) per electrode channel (Channel).

**Supplementary Table 1:** LME Results – Expectancy Step Effect in healthy Control Group.
***Excel File Name:*** Supplementary Table 1.xlsx
***Excel File Name in OSF Storage:*** LME_ExpectancyStep_CTR.xlsx

**Supplementary Table 2:** LME Results – Expectancy Step Effect in OFC Group
***Excel File Name:*** Supplementary Table 2.xlsx
***Excel File Name in OSF Storage:*** LME_ExpectancyStep_OFC.xlsx

**Supplementary Table 3:** LME Results – Expectancy Step Effect in LPFC Group
***Excel File Name:*** Supplementary Table 3.xlsx
***Excel File Name in OSF Storage:*** LME_ExpectancyStep_LPFC.xlsx

The following tables summarize results from LMEs with *expectancy step* (within-subject), group (between-subject), and their interaction, fitted separately for each pairwise comparison (OFC vs CTR, LPFC vs CTR, OFC vs LPFC). Each row corresponds to one EEG electrode (64 channels). Each table reports for the main effect of *expectancy step* the regression coefficient (column name: *b*_Distance), *t*-statistic (*t*_Distance), uncorrected *p*-value (p_Distance), and FDR-adjusted *p*-value (*p*_Distance_FDR); for the main effect of the group the regression coefficient (column name: *b*_Group), *t*-statistic (*t*_Group), uncorrected *p*-value (*p*_Group), and FDR-adjusted *p*-value (*p*_Group_FDR); and for the interaction *expectancy step X group* effect the regression coefficient (column name: *b*_Interaction), *t*-statistic (*t*_Interaction), uncorrected *p*-value (*p*_Interaction), and FDR-adjusted *p*-value (*p*_Interaction_FDR).

**Supplementary Table 4:** LME Results – Group Comparison (CTR vs. OFC)

***Excel File Name:*** Supplementary Table 4.xlsx
***Excel File Name in OSF Storage:*** LME_ExpectancyStep_CTR_vs_OFC.xlsx

**Supplementary Table 5:** LME Results – Group Comparison (CTR vs. LPFC)
***Excel File Name:*** Supplementary Table 5.xlsx
***Excel File Name in OSF Storage:*** LME_ExpectancyStep_CTR_vs_LPFC.xlsx

**Supplementary Table 6:** LME Results – Group Comparison (OFC vs. LPFC)
***Excel File Name:*** Supplementary Table 6.xlsx
***Excel File Name:*** LME_ExpectancyStep_OFC_vs_LPFC.xlsx

### NEUROPSYCHOLOGICAL TEST MEASURES:

**Supplementary Table 7. Demographics and neuropsychological test performance. Measures per participant group and statistical comparisons (CTR vs. OFC and CTR vs. LPFC).**

| **Demographics** | **CTR** | **SD** | **OFC** | **SD** | **LPFC** | **SD** | ***F* Value vs. OFC** | ***p* Value vs. OFC** | ***F* Value vs. LPFC** | ***p* Value vs. LPFC** |
|---|---|---|---|---|---|---|---|---|---|---|
| N | 15 | | 12 | | 10 | | | | | |
| Gender (females: males) | 10:5 | | 8:4 | | 6:4 | | | | | |
| Age years (range) | 48.5 (34-66) | 10.1 | 47.9 (27-61) | 11.7 | 40.9 (29-65) | 11.8 | 0.022 | 0.88 | 3.00 | 0.10 |
| Education years (range) | 16.5 (13-21) | 2.2 | 15 (9-21) | 3.1 | 15.9 (12-20) | 2.4 | 2.34 | 0.14 | 0.48 | 0.50 |
| **Neuropsychological tests** | | | | | | | | | | |
| Total IQ | 116.9 | 8.7 | 112.2 | 8.5 | 111.8 | 15.1 | 2.05 | 0.16 | 1.17 | 0.29 |
| Digit Span Total | 15.5 | 2.4 | 15 | 3.8 | 16.2 | 4.6 | 0.20 | 0.66 | 0.27 | 0.64 |
| Digit Span - Forward | 8.5 | 1.4 | 8.8 | 2.1 | 9.7 | 2.7 | 0.19 | 0.67 | 2.29 | 0.14 |
| Digit Span - Backward | 6.7 | 1.5 | 6.3 | 2.2 | 6.5 | 2.1 | 0.33 | 0.57 | 0.05 | 0.82 |
| Color-Word Interference Test (CWIT) | | | | | | | ***U* Value** | | ***U* Value** | |
| CWIT 1 – Color naming | 29.7 | 3.7 | 30.3 | 4.4 | 30.5 | 78.0 | 95.00 | 0.83 | 77.00 | 0.94 |
| CWIT 2 – Word reading | 21.9 | 2.8 | 22.1 | 3.7 | 22.3 | 5.1 | 84.00 | 0.79 | 75.50 | 1.00 |
| CWIT 3 – Inhibition | 51.6 | 9.0 | 52.3 | 11 | 55.4 | 17.5 | 90.00 | 1.00 | 87.00 | 0.50 |
| CWIT 4 – Inhibition/switching | 57.4 | 11.3 | 60.8 | 18.3 | 57.6 | 12.4 | 89.50 | 0.98 | 78.50 | 0.85 |

Comparison of the age, education, IQ (Wechsler Abbreviated Scale of Intelligence; WASI) and Digit Span Test (Wechsler Adult Intelligence Scale 3rd Edition; WAIS-III) between groups (One-Way ANOVA). Comparison of the non-normally distributed test scores, and the Colour-Word Interference Test (CWIT; Delis-Kaplan Executive Function System) between groups (non-parametric independent samples Mann-Whitney U test). Values are mean raw scores, with standard deviation (SD). IQ is reported as standard scores. CTR, healthy control group; LPFC, lateral prefrontal cortex lesion group; OFC, orbitofrontal lesion group; ns, not significant.

**INDIVIDUAL FRONTAL LESION MAPPING:**

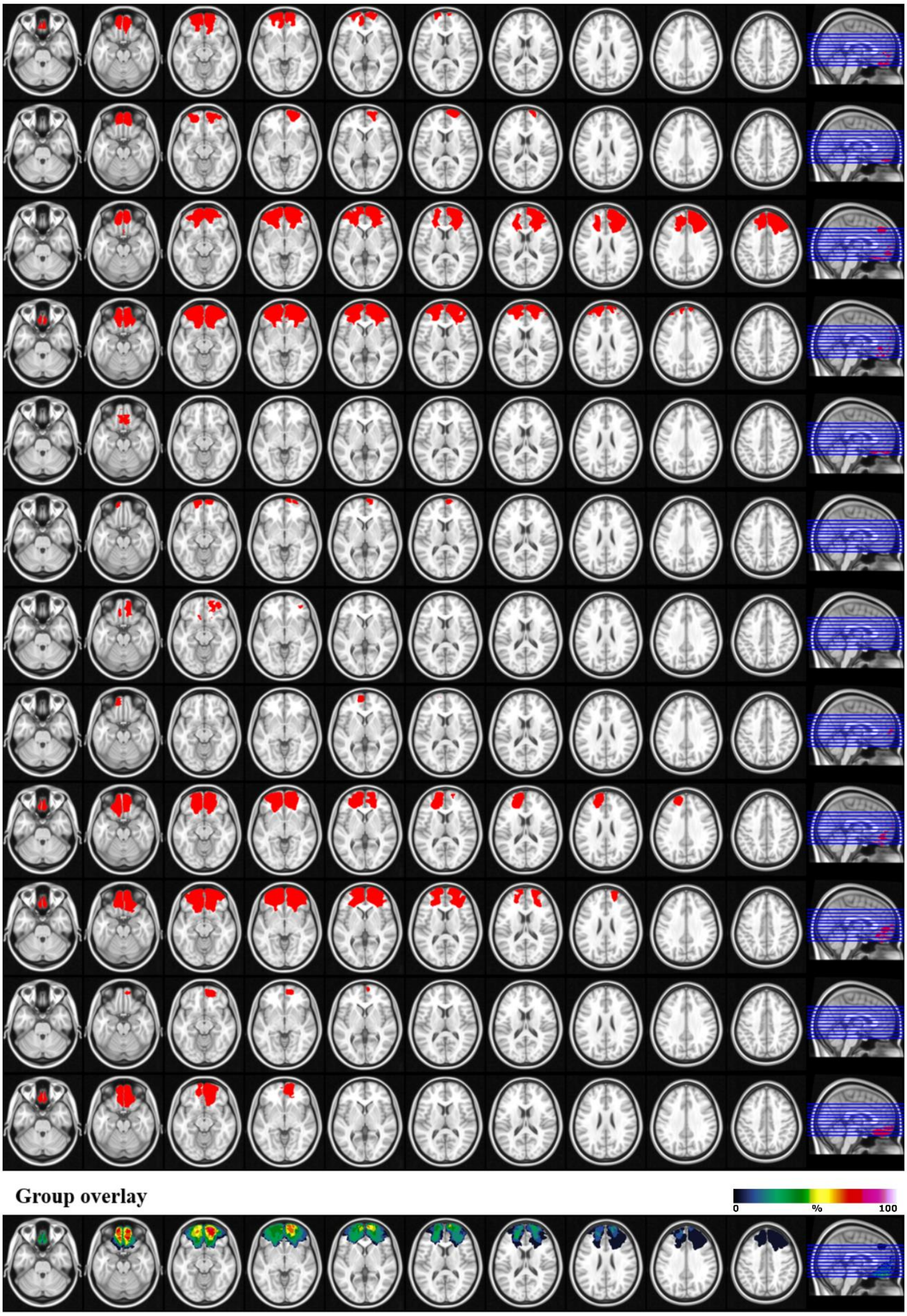


**Supplementary Figure 1. Lesion reconstructions for the orbitofrontal cortex (OFC) group.** Individual patients (1-12) and group overlay (bottom row). The colour code for the group overlay indicates the number of patients with damaged tissue in that area. Cortical lesion volume was concentrated in Brodmann areas 10, 11, 47, and 46, accounting for 87.1% of OFC lesion volume in the right hemisphere and 74.8% in the left (hemisphere-averaged ≈ 81.0%). Note: Each BA percentage is computed independently as the fraction of lesion volume overlapping that BA's atlas mask. Because the atlas masks overlap, the percentages do not sum to 100% (totals can exceed 100%).

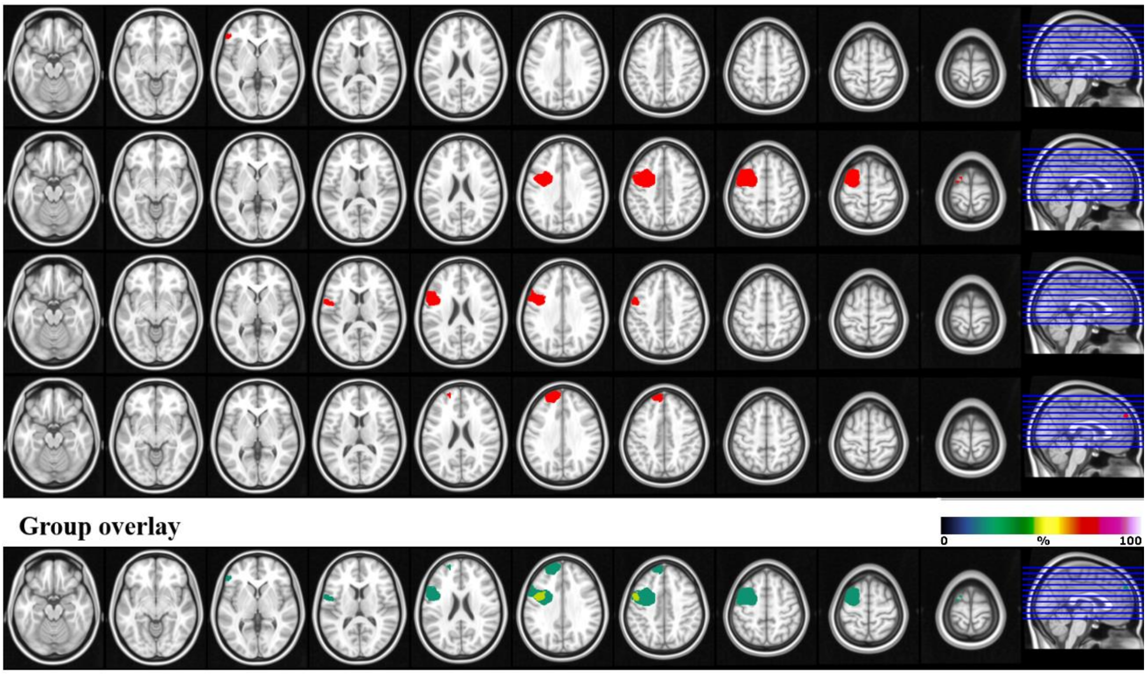


**Supplementary Figure 2. Lesion reconstructions for the left lateral prefrontal cortex (PFC) group.** Individual patients (1-4) and group overlay (bottom row). The colour code for the group overlay indicates the number of patients with damaged tissue in that area. Cortical lesion volume was concentrated in Brodmann areas 44, 6, and 9, accounting for 14.5% of the left-hemisphere LPFC lesion volume. Note: Each BA percentage is computed independently as the fraction of lesion volume overlapping that BA's atlas mask. Because the atlas masks overlap, the percentages do not sum to 100% (totals can exceed 100%).

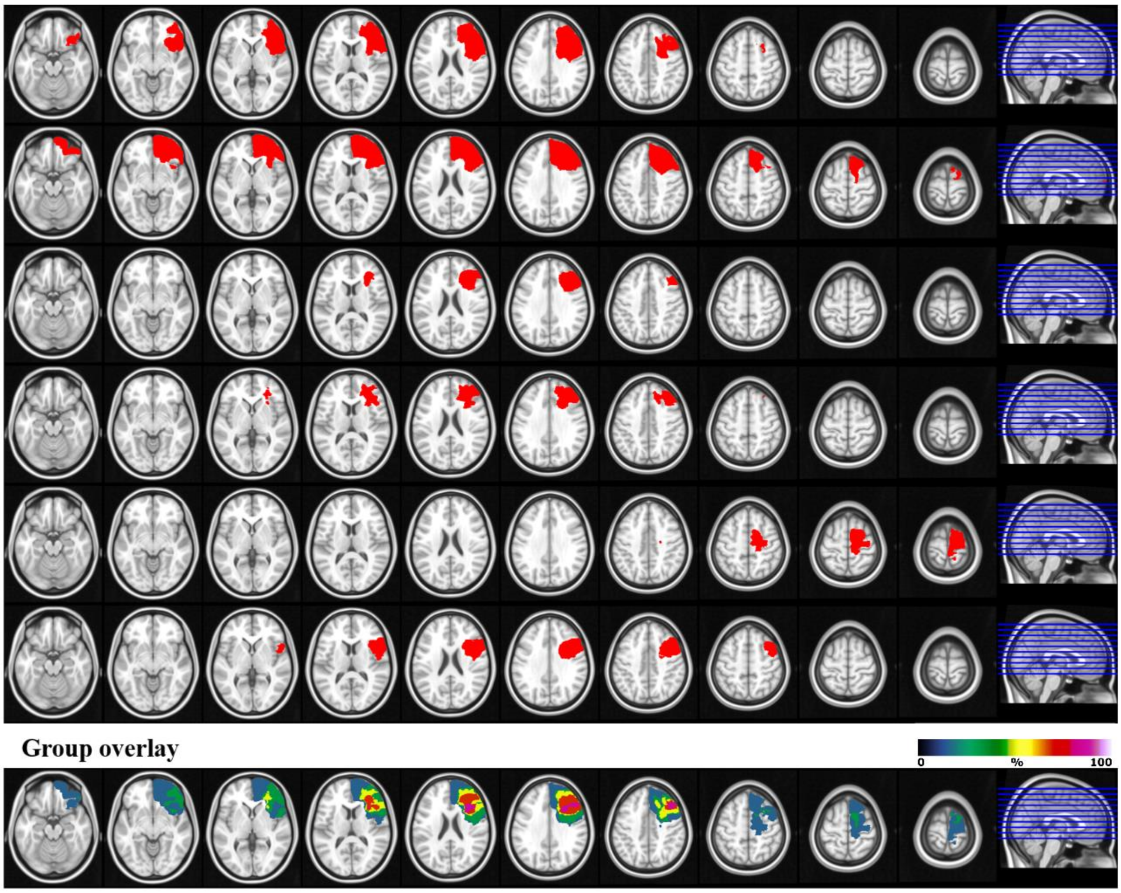


**Supplementary Figure 3. Lesion reconstructions for the right lateral prefrontal cortex (PFC) group.** Individual patients (1-6) and group overlay (bottom row). The colour code for the group overlay indicates the number of patients with damaged tissue in that area. Cortical lesion volume was concentrated in Brodmann areas 44, 46, 45, 47, 32, and 9 (29.6%, 25.8%, 25.1%, 16.5%, 15.2%, 15.1% respectively), accounting for 127.2% of right-hemisphere LPFC lesion volume. Note: Each BA percentage is computed independently as the fraction of lesion volume overlapping that BA's atlas mask. Because the atlas masks overlap, the percentages do not sum to 100% (totals can exceed 100%).

### <u>INTRACRANIAL STUDY</u>

#### EVENT-RELATED SEEG RESPONSES

From the intracranial EEG cohort, we extracted trial-wise event related responses (ERPs) from OFC (n = 90) and LPFC (n = 112) electrodes. To isolate a CNV-like slow potential, we subtracted the channel-wise mean evoked response. Because bipolar SEEG can invert polarity, we aligned polarity by enforcing that, for each channel, the 0-600 ms mean in the early steps condition (expectancy step = 2) was positive relative to the late step condition (expectancy step = 4 and 5). We then computed, for each trial, the 0-600 ms mean amplitude. We modeled these amplitudes with a single pooled linear mixed-effects model (LMEM) including region, *Amplitude ~ Expectancy Step × ROI + (1|Participant) + (1|Participantt: Electrode),* where *expectancy step* is the fixed effect of interest, ROI (OFC and LPFC) captures regional differences, and expectancy step X ROI interaction tests whether expectancy modulation differs between OFC and LPFC. Random intercepts for participant and electrode nested within participant account for repeated measures and channel-level variability. To control for biases, we repeated the entire pipeline in a permutation loop (N = 1000), permuting condition labels, re-aligning channel polarities, and refitting the LMEM on each surrogate. Surrogate fixed-effects *t*-values were used to build a null distribution of *t*-values. The resulting distribution was used to compute the *p*-values for the observed effect. This analysis showed that the mean evoked response encodes expectancy in both OFC ($p$ = 0.005) and LPFC ($p$ = 0.001).

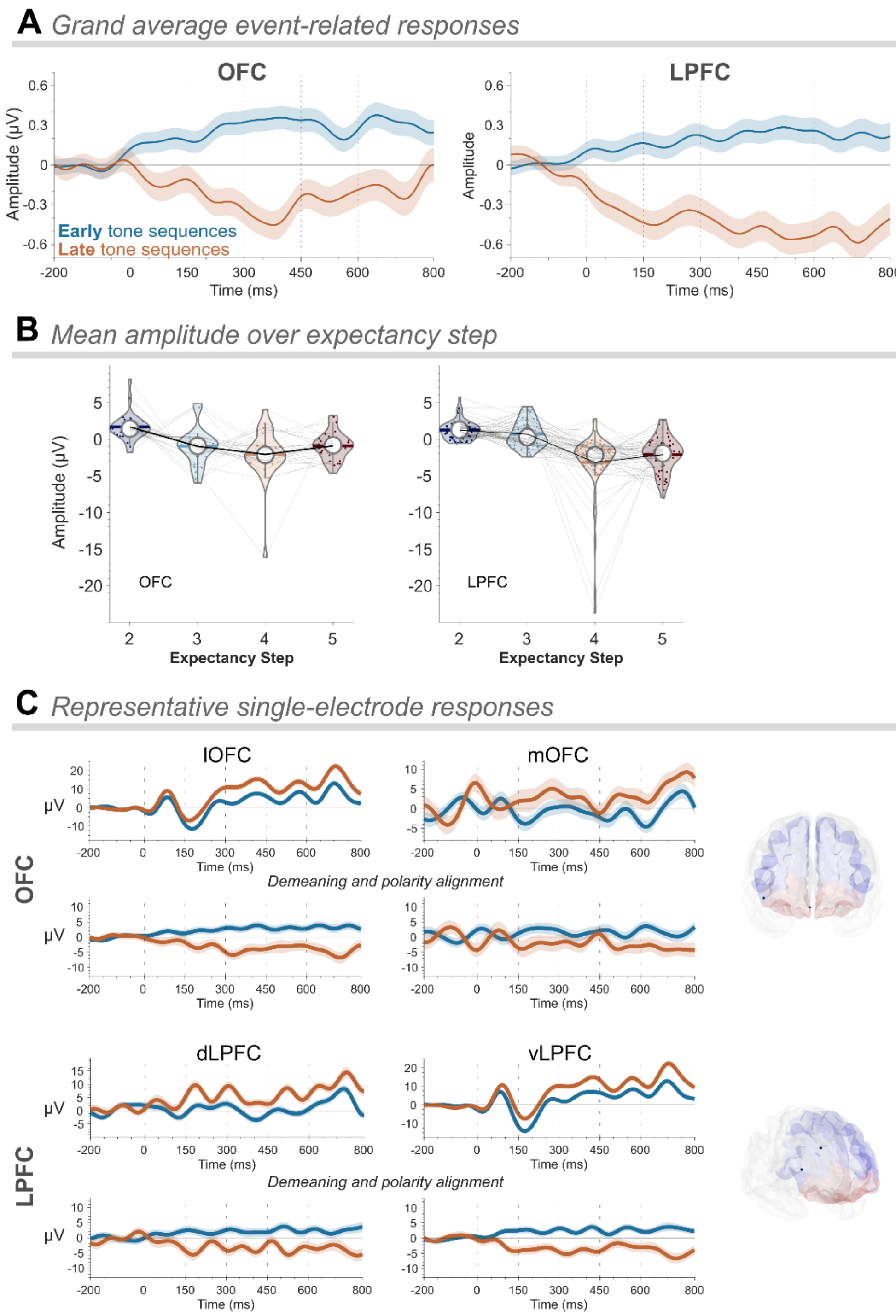


**Supplementary Figure 4. Intracerebral event-related responses in orbital, ventrolateral, and dorsolateral PFC are modulated by deviance expectancy.** (**A**) *Grand-average event-related responses.* Grand average waveforms recorded from orbitofrontal cortex (OFC; left, n = 90 electrodes) and lateral PFC (LPFC; right, n = 112 electrodes). Early tone sequences (*expectancy step* = 2) are shown in blue; late sequences (*expectancy steps* = 4 and 5) in brown. Vertical lines denote tone onsets. (**B**) *Mean amplitude over expectancy step.* Per-electrode mean amplitude over *expectancy step* (2–5) for OFC (left) and LPFC (right). Violin plots show the distribution across channels; points are electrode means; lines connect the same electrode across steps.

(**C**) *Representative single-electrode responses.* Event-related waveforms elicited by *early* (blue; step = 2) and late (brown; steps = 4+5) tone sequences recorded from dorsolateral PFC (dLPFC), ventrolateral PFC (vLPFC), medial OFC (mOFC), and lateral OFC (lOFC). Top rows show waveforms before demeaning and polarity alignment; bottom rows show the same electrodes after demeaning and polarity alignment.

### EPILEPTIC PATIENTS DEMOGRAPHICS AND CLINICAL CHARACTERISTICS:

**Supplementary Table 8. Patient demographics and clinical characteristics.**

| Patient | Sex | Age (y) | Education (y) | Implantation (hemisphere) | Epilepsy onset age (y) | Epilepsy duration (y) | Epileptogenic zone | MRI findings | ROI coverage (electrodes) | Other |
|---|---|---|---|---|---|---|---|---|---|---|
| 1 | M | 52 | N/A | R | 8 | 44 | Middle frontal sulcus | Small right frontal dysplasia (MFG) | OFC/LPFC 18/26 | |
| 2 | F | 50 | 16 | R/L | N/A | N/A | Left hippocampus | Tissue resection in right temporal lobe (MTL; 4 y prior) | OFC/LPFC 9/11 | |
| 3 | M | 33 | 11 | L | 13 | 20 | | None | OFC/LPFC 17/19 | |
| 4 | M | 32 | 17 | R | 19 | 13 | Ventral parietal cortex | Right ventral parietal lesion | OFC/LPFC 25/10 | |
| 5 | M | 48 | 14 | R | 1 | 47 | | None | OFC/LPFC 3/23 | |
| 6 | M | 39 | 17 | R | 16 | 23 | Temporal basal | None | OFC/LPFC 7/6 | |
| 7 | F | 41 | 13 | R | 28 | 13 | Posterior superior temporal sulcus | Right hippocampus | OFC/LPFC 11/17 | Dyslexia; migraine; reduced hearing |

The table lists, for each patient with epilepsy, demographics (sex, age, and education), the hemisphere where the electrodes were implanted, the age of onset of epileptic seizures, years since onset of epilepsy, the epileptogenic zone, Magnetic Resonance Imaging (MRI) findings, frontal Region of Interest (ROI) electrode coverage (OFC, orbitofrontal cortex; LPFC, lateral prefrontal cortex), and other clinical characteristics. y, year; R, right; L, left; MTL, medial temporal lobe; MFG, middle frontal gyrus.

### TASK DESIGN

**Supplementary Table 9. Expectancy indices used in the task design.**

| *Index* | *What it represents* | *Formula* | *Notes / Usage* |
|---|---|---|---|
| **A priori distance** | Design distribution of distances between consecutive deviants | $P$ ($L$ = 2, 3, 4, 5) {25%, 30%, 10%, 35%} | Describes stimulus statistics (chunk lengths). Not the probability that the next sequence is deviant. |
| **Conditional ("hazard")** | Probability the next sequence is deviant, given no deviant has occurred yet | $P$ (dev at $s$ \| $L \geq s$) {25%, 40%, 22%, 100%} | What an ideal observer would use to update expectancy online. |

| | | | |
|---|---|---|---|
| **Monotonic step-index (heuristic)** | Ordinal proxy "more elapsed standards ⇒ higher expectancy" | E($s$) = ($s$ − 1) / 4 × 100% {25%, 50%, 75%, 100%} | Not an objective probability; it reflects monotonic rise across steps. |

$s$ = **expectancy step** (position of the next sequence since the last deviant tone sequence); $L$ = **distance** (number of tone sequences / chunk length) between consecutive deviant tone sequences. $s = L$ only when the deviant occurs.
